# Unraveling the Intricate Cargo-BBSome Coupling Mechanism at the Ciliary Tip

**DOI:** 10.1101/2022.11.02.514950

**Authors:** Yan-Xia Liu, Wen-Juan Li, Rui-Kai Zhang, Sheng-Nan Sun, Zhen-Chuan Fan

**Affiliations:** State Key Laboratory of Food Nutrition and Safety, Institute of Health Biotechnology, Tianjin University of Science and Technology, Tianjin 300457, China

**Keywords:** Cilia, BBSome, ARL13, BBS3, intraflagellar transport, Bardet-Biedl syndrome

## Abstract

Certain ciliary transmembrane and membrane-tethered signaling proteins migrate from the ciliary tip to base via retrograde intraflagellar transport (IFT), essential for maintaining their ciliary dynamics to enable cells to sense and transduce extracellular stimuli inside the cell. During this process, the BBSome functions as an adaptor between retrograde IFT trains and these signaling protein cargoes. The Arf-like 13 (ARL13) small GTPase resembles ARL6/BBS3 in facilitating these signaling cargoes to couple with the BBSome at the ciliary tip prior to loading onto retrograde IFT trains for transporting towards the ciliary base, while the molecular basis for how this intricate coupling event happens remains elusive. Here, we report that *Chlamydomonas* ARL13 only in a GTP-bound form (ARL13^GTP^) anchors to the membrane for diffusing into cilia. Upon entering cilia, ARL13 undergoes GTPase cycle for shuttling between the ciliary membrane (ARL13^GTP^) and matrix (ARL13^GDP^). To achieve this goal, the ciliary membrane-anchored BBS3^GTP^ binds and activates the ciliary matrix-residing ARL13^GDP^ as an ARL13 guanine nucleotide exchange factor. At the ciliary tip, ARL13^GTP^ binds and recruits the ciliary matrix-residing and post-remodeled BBSome as an ARL13 effector to anchor to the ciliary membrane. This makes the BBSome spatiotemporally become available for the ciliary membrane-tethered phospholipase D (PLD) to couple with. Afterward, ARL13^GTP^ hydrolyzes GTP for releasing the PLD-laden BBSome to load onto retrograde IFT trains. According to this model, hedgehog signaling defects associated with *ARL13b* and *BBS3* mutations in humans could be satisfactorily explained, providing us a mechanistic understanding behind BBSome-cargo coupling required for proper ciliary signaling.

**Significance statement:** Certain signaling proteins export in a BBSome-dependent manner from cilia via retrograde IFT. As IFT cargoes, their coupling with the BBSome is mediated by ARL13 and BBS3. Here, we report that ARL13 in a GTP-bound state binds the membrane for diffusing into cilia followed by running GTPase cycle with BBS3 acting as an ARL13-specific guanine nucleotide exchange factor. ARL13^GTP^ recruits the IFT-shed BBSome as its effector to anchor to the membrane of the ciliary tip, making it spatiotemporally available for the ciliary membrane-tethered PLD to bind. Afterward, ARL13 GTP hydrolysis releases the PLD-laden BBSome for retrograde IFT train loading. ARL13 and BBS3 facilitate the cargo-BBSome coupling in such a way providing a regulatory mechanism for signaling protein export from cilia.

## Introduction

Ciliary transmembrane (e.g. ion channels and G protein-coupled receptors, GPCRs) and membrane-tethered (e.g. phospholipase D, PLD) signaling proteins sense and transduce the extracellular stimuli into the cell (1). To function properly, they cycle through cilia as intraflagellar transport (IFT) cargoes (2). During this process, the octameric BBSome bridges them to IFT trains composed of complexes IFT-A and -B (subdivided into IFT-B1 and -B2) (3–5). Therefore, defects in BBSome coupling with cargoes or IFT trains disbalance these signaling proteins in the ciliary membrane, leading to Bardet-Biedl syndrome (BBS) (6, 7). Interestingly, the mutated Arf-like 3 (ARL3) and ARL13b small GTPases cause Joubert syndrome (JBTS) with symptoms overlapped with BBS (8–12). We have known that ARL13b acts as a guanine nucleotide exchange factor (GEF) for ARL3 in cilia (13–17). The active ARL3^GTP^ binds the carrier proteins PDE6D and UNC119A/B for releasing their cytoplasmic lipidated protein cargoes to anchor to the ciliary membrane (13, 18–20). Thus, mutations in *ARL3* and *ARL13b* were implicated to cause JBTS by disabling signaling protein import into cilia (17, 19, 21–27).

ARL3 and ARL13b also mediate signaling protein export from cilia in a BBSome-dependent manner but via distinct mechanisms. As reflected in *Chlamydomonas reinhardtii,* ARL3 with the aid of Rab-like 2 (RABL2) small GTPase acting as an ARL3 GEF recruits the PLD-laden BBSome as an ARL3 effector to diffuse through the transition zone (TZ) for ciliary retrieval (28–30). ARL13, the ARL13b orthologue of humans, instead enables PLD coupling with the BBSome for exporting from cilia via the IFT/BBS system (11). Since many GPCRs like somatostatin receptor 3 and GPR161 and smoothened (Smo) of hedgehog signaling (Hh) pathway couple with the BBSome for loading onto retrograde IFT trains for ciliary retrieval (31–37), roles of ARL13 and ARL3 in PLD coupling with the BBSome (ARL13) and PLD-laden BBSome diffusion through the TZ for ciliary retrieval (ARL3), respectively, well explain their buildup in certain Arl13b and Arl3 mutant cilia of rodent cells (14, 38–40). If these hold true in humans, mutations in *ARL3* and *ARL13b* genes cause BBS could be satisfactorily explained (11, 28).

The BBSome sheds from anterograde IFT at the ciliary tip and undergoes disassembly followed by reassembly, a process called BBSome remodeling (34, 41–47). At the ciliary tip of rodent cells, the small GTPase Rabl4/Ift27 activates Arl6/Bbs3 by acting as a Bbs3 GEF, a process indispensable for the GPCR-laden BBSome likely as a Bbs3 effector to load onto retrograde IFT trains for ciliary retrieval (34, 48). As reflected in *C. reinhardtii,* IFT27 promotes BBSome ciliary tip reassembly, making the post-remodeled BBSome available for retrograde IFT loading for ciliary retrieval (41). BBS3 cannot facilitate BBSome loading onto retrograde IFT trains but resembles ARL13 in enabling PLD coupling with the BBSome at the ciliary tip (11, 49). This coupling event can happen even when BBS3 GTPase cycle is blocked by providing BBS3 of a GTP-locked state in cilia (49). Besides these, the ciliary tip cargo-BBSome coupling mechanism unfortunately has remained uncertain.

By using *C. reinhardtii* as a model organism, which supports unbiased biochemical analysis of isolated cilia and the cell body (50, 51), we herein report that ARL13^GTP^ anchors to the membrane at the basal body for diffusing into cilia prior to its GTPase cycle with BBS3^GTP^ acting as an ARL13 GEF. ARL13^GTP^ binds and anchors the post-remodeled BBSome as its effector to the membrane of the ciliary tip, enabling the ciliary membrane-tethered PLD to couple with. ARL13^GTP^ then hydrolyzes GTP to release the PLD-laden BBSome for retrograde IFT train loading. Unraveling this intricate ciliary tip cargo-BBSome coupling mechanism thus provides us novel insights into the molecular basis of BBS disorders in humans.

## Results

### ARL13^GTP^ anchors to the membrane for diffusing into cilia followed by a conversion to ARL13^GDP^

Either GTP- or GDP-bound state and the N-terminal amphipathic helix are two factors required for Arf-like GTPases to anchor to the membrane (28, 49, 52). To dissect how they affect ARL13 anchoring to the membrane, we expressed ARL13 and its variants harboring the mutation G73L, T33N, or the N-terminal 15 residue deletion (ΔN15) combined with G73L, T33N, or none of them in bacteria (Fig. 1*A*). G73L and T33N were introduced in ARL13 or ARL13ΔN15 as they supposedly render ARL13 active (GTP-bound) or inactive (GDP-bound or nucleotide-free), respectively (*SI Appendix,* Fig. S1*A*). ARL13 preloaded with GTPγS but not GDP was bound to liposomes, identifying ARL13^GTP^ but not ARL13^GDP^ interacts with liposomes (Fig. *1B*). Consistently, ARL13G73L was bound to liposomes and ARL13T33N did not do so (Fig. 1*B*). Besides, ARL13ΔN15 failed to bind liposomes and remained unbound to liposome even when G73L was introduced (Fig. 1*B*). Thus, the N-terminal amphipathic helix and GTP-bound state are two prerequisites for ARL13 to anchor to the membrane. To dissect how membrane anchoring affects ARL13 ciliary entry, we expressed ARL13 and its variants fused at their C-terminus to HA-YFP in the ARL13-null CLiP mutant that we named *arl13-356* (LMJ.RY0402.181356) (11). The resulting six strains expressed the tagged ARL13 and its variants at the parental CC-5325 ARL13 level (Fig. 1*B* and *SI Appendix,* Fig. S1*B*) (11). Notably, ARL13G73L-HA-YFP rather than the T33N variant resembled the endogenous ARL13 and ARL13-HA-YFP in residing in cilia, while ARL13ΔN15-HA-YFP and its G73L and T33N versions all failed to enter cilia, identifying ARL13^GTP^ enters cilia via anchoring to the membrane under physiological conditions (Fig. 1*B*).

**Figure 1.**
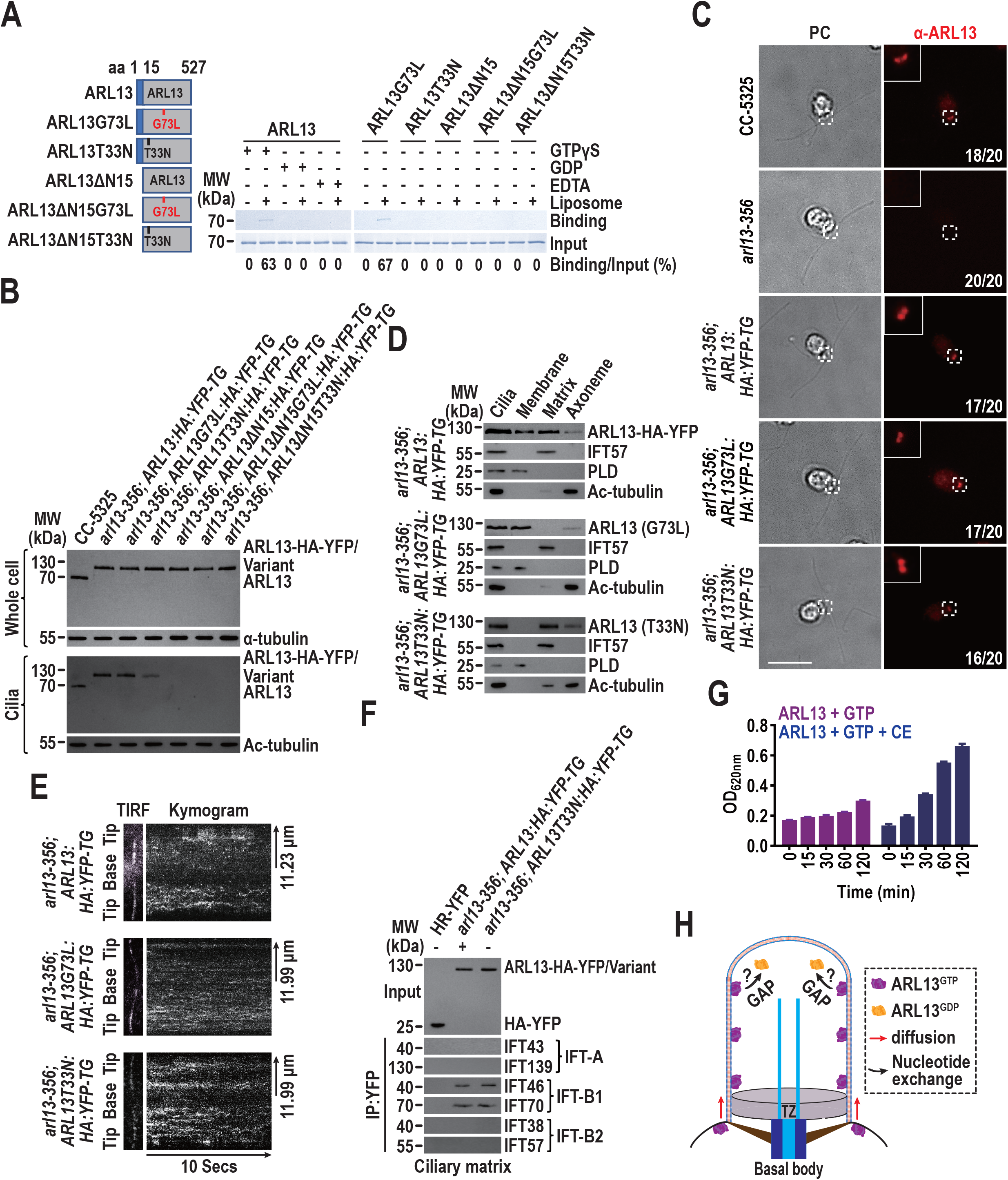
ARL13^GTP^ anchors to the membrane for diffusing into cilia followed by a conversion to ARL13^GDP^. (*A*). Schematic presentation of bacterially expressed ARL13 and its variants (shown on the left). ΔN15 stands for the N-terminal 15 amino acids of ARL13 deleted. Immunoblots of liposome flotation captured ARL13 preloaded with GTPγS or GDP (shown in the middle) and its variants (shown on the right) with α-ARL13. ARL13 or its variants along was used as an input for evaluating their binding/input ratio shown as percentile. (*B*). Immunoblots of whole cell samples and cilia of cells indicated on the top probed with α-ARL13. Alpha-tubulin and acetylated (Ac)-tubulin were used to adjust the loading of whole cell samples and cilia, respectively. MW: molecular weight. (*C*). Cells shown on the left stained with α-ARL13 (red). Phase contrast (PC) images of cells were shown. Insets are 9 times enlargements of the basal bodies indicated with a white box. Scale bar: 10 μm. Cell numbers out of 20 cells are listed for representing cells that are ARL13 positive or negative at the basal bodes. (*D*). Immunoblots of ciliary fractions of cells indicated on the left probed with α-ARL13, α-IFT57 (ciliary matrix marker), α-PLD (ciliary membrane marker) and Ac-tubulin (axoneme marker). MW: molecular weight. (*E*). Representative TIRF images and corresponding kymograms of cells indicated on the left (Video 1-3, 15 fps). The time and transport lengths are indicated on the bottom and right, respectively. The ciliary base (base) and tip (tip) were shown. (*F*). Immunoblots of α-YFP-captured proteins from the ciliary matrix of cells indicated on the top (in the presence of GDP for *arl13; ARL3:HA:YFP-TG* ciliary matrix) probed for the IFT-A subunits IFT43 and IFT139, the IFT-B1 subunits IFT46 and IFT70, and the IFT-B2 subunits IFT38 and IFT57. Input was quantified with α-YFP by immunoblotting. MW: molecular weight. (*G*). GTP hydrolysis of ARL13 in the absence and presence of ciliary extracts (CE) isolated from CC-5325 cells. ARL13 shows low intrinsic GTPase activity. The presence of CE elevates the GTPase activity of ARL13 remarkably. (*H*) Schematic presentation of how ARL13^GTP^ binds the membrane for diffusing into cilia prior to converting to ARL3^GDP^ for shifting to reside in the ciliary matrix.

As reflected by ARL13-HA-YFP, ARL13 was enriched at the basal bodies and resided in the ciliary membrane, matrix, and axoneme (Fig. 1*C* and *D* and *SI Appendix,* Fig. S1*C*) (11, 53). It diffused in cilia with very occasional IFT movement hard to capture and distributed along the whole length of cilia (Fig. *1E* and Movie 1) (11). According to ARL13G73L-HA-YFP and the GTPγS-loaded ARL13, ARL13^GTP^ was concentrated at the basal bodies at normal level and resided in the ciliary membrane and axoneme but not matrix (Fig. 1*C* and *D* and *SI Appendix,* Fig. S1*C*). It was moved solely by diffusion and distributed along the whole length of cilia (Fig. 1*E* and Movie 2). ARL13T33N-HA-YFP and the GDP-loaded ARL13 recapitulated the endogenous ARL13 in positioning to the basal bodies, distributing in cilia, and moving in cilia and resided in the ciliary matrix and axoneme but not membrane (Fig. 1*C-E* and *SI Appendix,* Fig. S1*C* and Movie 3). Thus, ARL13 can associate with axoneme without relying on its nucleotide state, whereas nucleotide exchange enables ARL13 to shuttle between the ciliary membrane and matrix. Since ARL13 resides in both the ciliary membrane and matrix but dominantly as a soluble ciliary matrix protein (Fig. 1*D* and *SI Appendix,* Fig. S1*C*) (11), ARL13^GTP^, upon entering cilia, must have to convert to ARL13^GDP^. In view of the fact that the GDP-loaded ARL13-HA-YFP and the T33N variant both immunoprecipitated IFT-B1 but not IFT-A and -B2 in the ciliary matrix, it could be ARL13^GDP^ that performs processive IFT via interacting transiently with IFT-B1 (Fig. 1*F*) (11). Besides, ARL13 had low intrinsic GTP hydrolysis activity, while ciliary extracts dramatically elevated its GTPase activity, indicative of the existence of a yet-to-be-determined GTPase-activating protein (GAP) for ARL13 in cilia (Fig. 1*G*) (18). In sum, it is ARL13^GTP^ that anchors to the membrane for diffusing into cilia followed by a conversion to ARL13^GDP^ most likely in an ARL13 GAP-dependent manner (Fig. *2H*).

**Figure 2.**
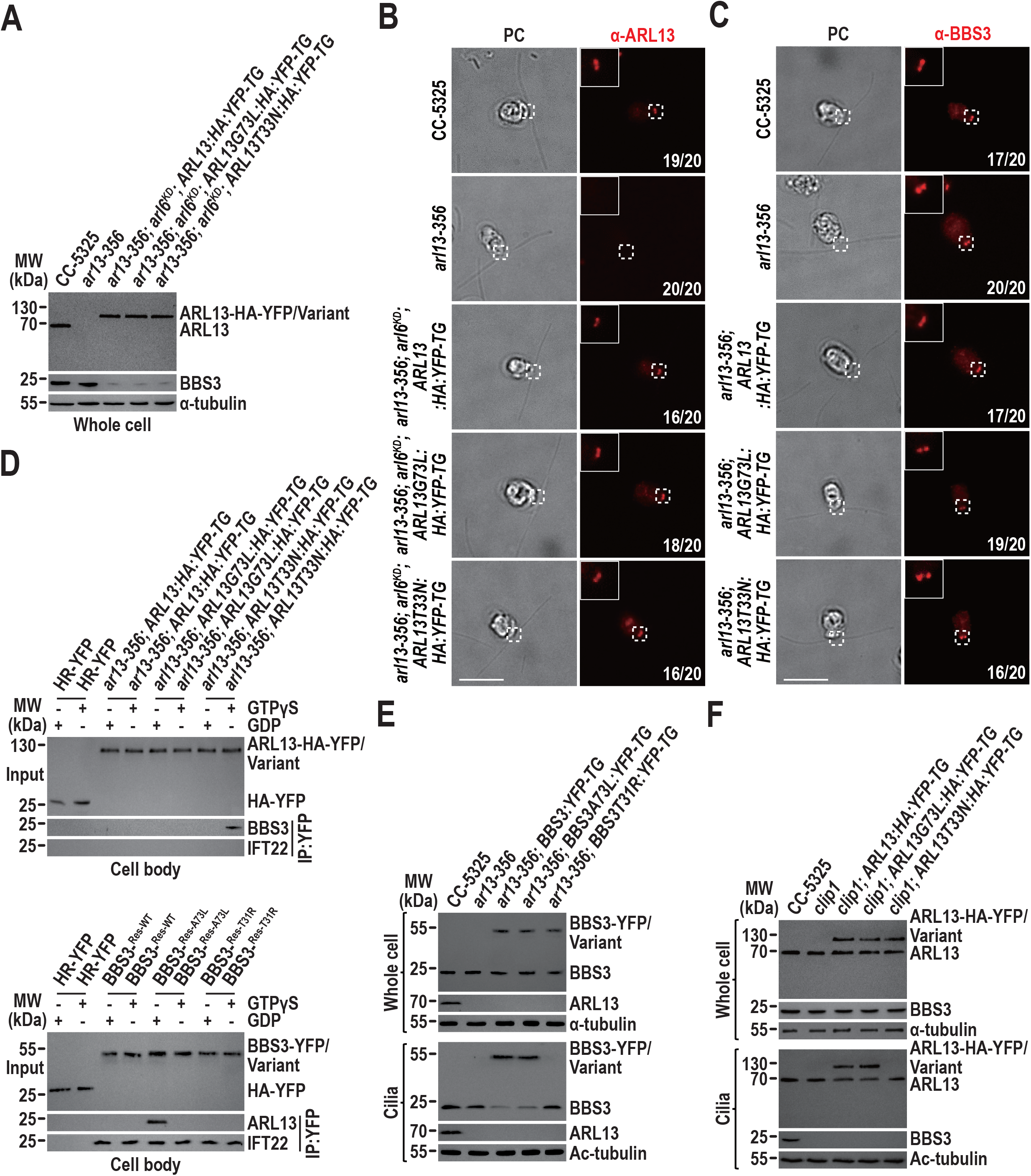
ARL13^GTP^ and BBS3^GTP^ traffic to the basal body for ciliary entry independently. (*A*). Immunoblots of whole cell samples indicated on the top probed with α-ARL13 and α-BBS3. Alphatubulin was used as a loading control. MW: molecular weight. (*B-C*). Cells shown on the left stained with α-ARL13 (red) (*B*) or α-BBS3 (red) (*C*). For both panels, PC images of cells were shown. Insets are 9 times enlargements of the basal bodies indicated with a white box. Scale bar: 10 μm. Cell numbers out of 20 cells are listed for representing cells that are ARL13 (*B*) or BBS3 (*C*) positive or negative at the basal bodes. Scale bar: 10 μm. (*D*). Immunoblots of α-YFP-captured proteins from cell body extracts of cells indicated on the top probed with α-ARL13, α-BBS3, and α-IFT22. Input was quantified with α-YFP by immunoblotting. Alpha-tubulin was used to adjust the loading. MW: molecular weight. The assays were performed in the presence of GTPγS or GDP. (*E-F).* Immunoblots of whole cell samples and cilia of cells indicated on the top probed with α-BBS3 and α-ARL13. Alpha-tubulin and acetylated (Ac)-tubulin were used to adjust the loading of whole cell samples and cilia, respectively. MW: molecular weight.

### ARL13^GTP^ and BBS3^GTP^ traffic to the basal bodies for ciliary entry independently

ARL13 can translocate from the cell body to the basal bodies in a nucleotide-independent manner as the G73L and T33N variants both can target to and reside at the basal bodies at the endogenous ARL13 level (Fig. 1*C*). BBS3 knockdown to as low as ~10% of its endogenous level did not alter the basal body contents of ARL13-HA-YFP and the G73L and T33N variants, excluding BBS3 from promoting ARL13 translocation to the basal bodies (Fig. 2*A* and *B*). Reciprocally, BBS3 did not require ARL13 for doing so as loss of ARL13 retained normal BBS3 basal body abundance (Fig. 2*C*). In cytoplasm, ARL13-HA-YFP failed to immunoprecipitate BBS3 in the presence of GTPγS or GDP, which render two GTPases GTP- or GDP-bound simultaneously; the T33N variant recovered BBS3 only when GTPγS was present to render BBS3 GTP-bound; and none of GTPγS and GDP conferred BBS3 to be recovered by the G73L variant, identifying BBS3^GTP^ interacts with ARL13^GDP^ (Fig. *2D*). BBS3 was found to bind RABL5/IFT22 to form IFT22/BBS3 in cytoplasm (54). Only when they both are GTP-bound, IFT22/BBS3 traffics to the basal bodies, where BBS3^GTP^ uncouples with IFT22^GTP^ before it can anchor to the membrane for diffusing into cilia (49, 54). Reflecting this fact, BBS3-YFP, BBS3A73L-YFP (GTP-bound), and BBS3T33N-YFP (GDP-bound) all immunoprecipitated IFT22 without depending on IFT22 nucleotide state in cytoplasm and only BBS3A73L-YFP recovered ARL13 simultaneously (Fig. 2*D*) (54). Therefore, BBS3^GTP^ binds IFT22^GTP^ for trafficking to the basal bodies, where it uncouples with IFT22^GTP^ for shifting to interact with ARL13^GDP^ (49, 54).

To discern how BBS3^GTP^ and ARL13 interplay for entering cilia from the basal bodies, we expressed BBS3-YFP, BBS3A73L-YFP, and BBS3T31R-YFP at the same level in *arl13-356* cells to generate *arl13-356; BBS3:YFP-TG, arl13-356; BBS3A73L:YFP-TG,* and *arl13-356; BBS3T31R:YFP-TG* strains (Fig. 2*E*) (49). Loss of ARL13 did not impede ciliary entry of the endogenous BBS3, BBS3-YFP, and BBS3A73L-YFP (Fig. *2E*). BBS3T31R-YFP failed to enter cilia as it is disabled to traffic to the basal bodies, revealing that BBS3^GTP^ does not require ARL13 for entering cilia from the basal body (Fig. 2*E*) (54). We next expressed ARL13-HA-YFP and its G73L and T33N variants at the same level in the *clip1* mutant to generate strains *clip1; ARL13:HA:YFP-TG*, *clip1; ARL13G73L:HA:YFP-TG*, and *clip1; ARL13T33N:HA:YFP-TG* (Fig. 2*F*). In *clip1* mutant, BBS3^GTP^ retains the ability to traffic to the basal body but somehow fails to enter cilia (49). Remarkedly, loss of BBS3 in cilia only disabled ARL13T33N-HA-YFP to enter cilia (Fig. 2*F*). Thus, ARL13^GDP^ rather than ARL13^GTP^ relies on BBS3^GTP^ for entering cilia from the basal bodies, revealing that ARL13^GDP^ diffuses into cilia via binding BBS3^GTP^ transiently (Fig. *1B*). Accordingly, the ARL13ΔN15T33N variant can traffic to the basal bodies but failed to enter cilia as loss of the N-terminal 15 amino acids disrupted ARL13^GDP^ binding to BBS3^GTP^ (Fig. 1*B* and *SI Appendix,* Fig. *S2A* and *B*).

### BBS3^GTP^ activates ARL13^GDP^ as an ARL13 GEF in cilia

ARL13 resided in the ciliary matrix in *clip1* cilia (Fig. 3*A*). Compared with CC-5325 cilia, ARL13 was completely lost in the ciliary membrane of *clip1* cilia, identifying a BBS3-dependent conversion of ARL13^GDP^ to ARL13^GTP^ must occur in *Chlamydomonas* cilia (Fig. 3*A*). ARL13^GDP^ requires BBS3 for converting to ARL13^GTP^ in cilia, indicative of BBS3 as an ARL13 GEF in cilia (Fig. 3*B*). To answer this question, we crossed the *bbs8* mutant with *arl13; ARL13:HA:YFP-TG, arl13; ARL13Q73L:HA:YFP-TG,* and *arl13; ARL13T30N:HA:YFP-TG* cells to generate strains *arl13; bbs8; ARL13:HA:YFP-TG, arl13; bbs8; ARL13Q73L:HA:YFP-TG,* and *arl13; bbs8; ARL13T30N:HA:YFP-TG* (*SI Appendix,* Fig. S3*A*) (55). By doing so, these strains lost the BBSome in cilia due to the lack of intact BBSomes caused by BBS8 knockout but contained BBS3 of normal abundance in cilia (*SI Appendix,* Fig. S3*A*) (28). In the presence of GTPγS, ARL13-HA-YFP and ARL13G73L-HA-YFP immunoprecipitated BBS3 and so did for the T33N variant (Fig. 3*C*). ARL13-HA-YFP and its G73L and T33N versions all failed to recover the GDP-loaded BBS3 (Fig. 3*C*). We next expressed BBS3-YFP and BBS3A73L-YFP at the same level in the *bbs8* mutant to generate *bbs8; BBS3:YFP-TG* and *bbs8; BBS3A73L:YFP-TG* strains, whose cilia contained BBS3-YFP or BBS3A73L-YFP, maintained normal ARL13, and lost the BBSome (*SI Appendix,* Fig. S3*B*) (49). BBS3-YFP pre-loaded with GTPγS and BBS3A73L-YFP immunoprecipitated ARL13 in cilia and the GDP addition only deprived BBS3-YFP of ARL13 recovery (Fig. *3D*). Besides, BBS3A73L-YFP and ARL13 cofractionated in *bbs8; BBS3A73L:YFP-TG* cilia at a molecular weight of ~111 kDa in the presence of GTPγS or GDP (Fig. 3*E*). Thus, BBS3^GTP^ can transiently bind ARL13 directly in cilia in an ARL13 nucleotide state- and BBSome-independent manner.

**Figure 3.**
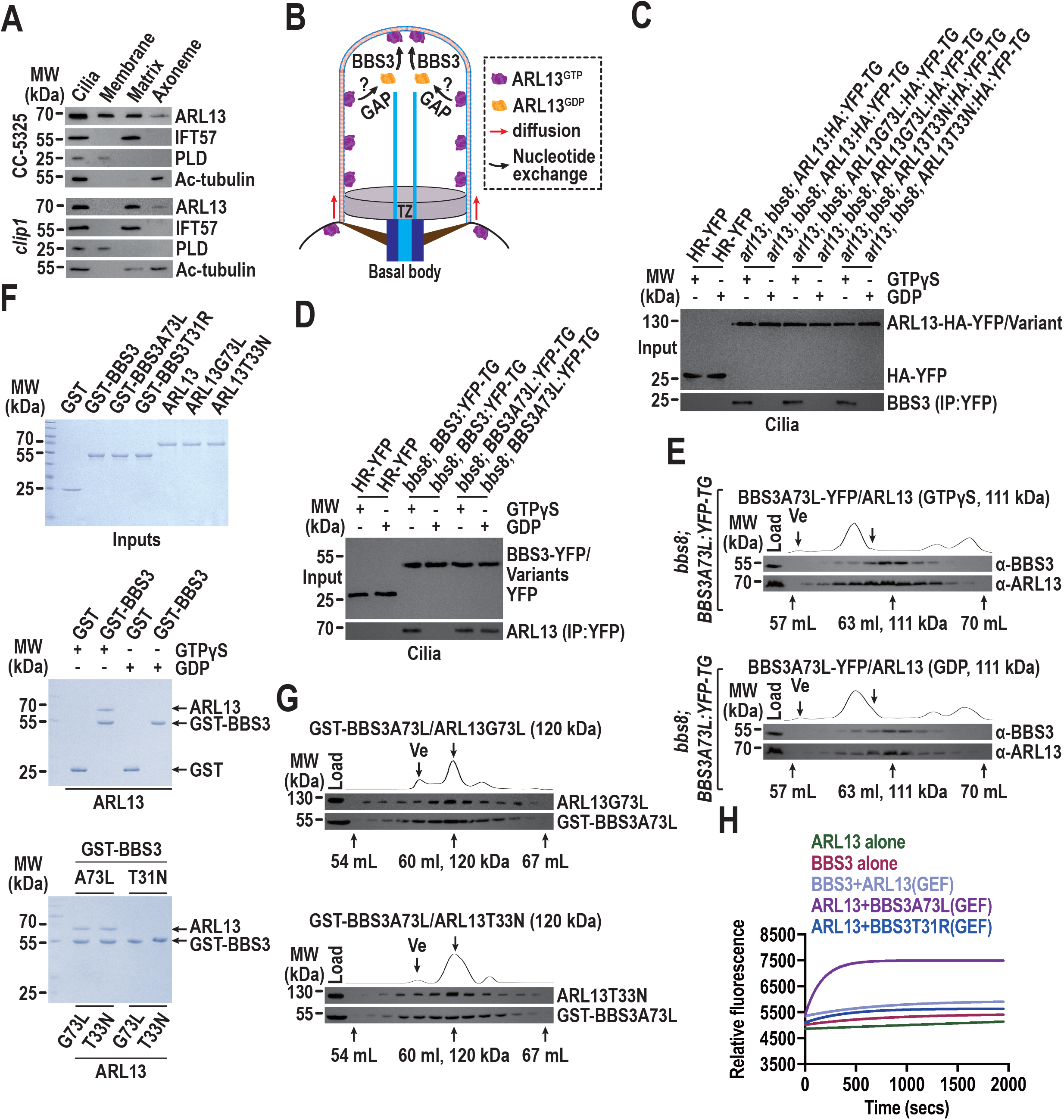
BBS3^GTP^ activates ARL13^GDP^ as an ARL13 GEF in cilia. (*A*). Immunoblots of ciliary fractions of cells indicated on the left probed with α-ARL13, α-IFT57 (ciliary matrix marker), α-PLD (ciliary membrane marker) and Ac-tubulin (axoneme marker). (*B*). Schematic presentation of ARL13 GTPase cycle in cilia. ARL13^GTP^ binds the membrane for diffusing into cilia, converts to ARL3^GDP^ most likely in a GAP-dependent manner for shifting to reside in the ciliary matrix, and ARL13^GDP^ converts back to ARL13^GTP^ with the aid of BBS3 for anchoring to the ciliary membrane. (*C-D*). Immunoblots of α-YFP-captured proteins from cilia of cells indicated on the top in the presence of GTPγS or GDP probed for BBS3 (*C*) or ARL13 (*D*). Input was quantified with α-YFP by immunoblotting. MW: molecular weight. (*E*). Cilia of cells indicated on the left fractionated by size exclusion chromatography (S200 sizing column) and probed with α-BBS3 and α-ARL13 by immunoblotting. Relative protein concentration of the fractions between elution volume of 57 ml and 70 ml is shown as absorbance at 280 nm. The excluded volume (Ve) and the elution volume of 63 ml (111 kDa) in the presence of GTPγS and GDP are indicated under the immunoblots. (*F*). GST, GST-BBS3, GST-BBS3A73L, GST-BBS3T31R, ARL13, ARL13G73L, and ARL13T33N resolved by SDS-PAGE and visualized by Coomassie staining (Upper). GST and GST-BBS3 mixed with ARL13 in the presence of GTPγS or GDP and complexes recovered on glutathione beads resolved by SDS-PAGE and visualized by Coomassie staining (Middle). GST-BBS3A73L, GST-BBS3T31R, ARL13G73L, and ARL13T33N mixed to form four combinations and complexes recovered on glutathione beads resolved by SDS-PAGE and visualized by Coomassie staining (Bottom). (*G*). GST-BBS3A73L/ARL13G73L and GST-BBS3A73L/ARL13T33N recovered on glutathione beads fractionated by size exclusion chromatography (S200 sizing column) and probed with α-BBS3 and α-ARL13 by immunoblotting. Relative protein concentration of the fractions between elution volume of 54 and 67 ml is shown as absorbance at 280 nm. The excluded volume (Ve) and the elution volume of 60 ml (120 kDa) are indicated under the immunoblots. (*H*). Mant-GTP association measurement for determining GEF activity of BBS3 on ARL13. Fluorescence intensity of mant-GTP (0.75 μm) was plotted against the recording time (s) for a combination of 2 μm ARL13, BBS3, BBS3A73L, or BBS3T31R alone or in the presence of 0.5 μm BBS3, BBS3A73L, or BBS3T31R (GEF activity on ARL13) or ARL13 (GEF activity on BBS3). Data are averaged from three replicates. The experiment was performed three times, and one representative is shown.

We next generated bacterially expressed N-terminal GST-tagged BBS3 (GST-BBS3), ARL13, and their variants (Fig. 3*F*). ARL13 failed to bind GST but was bound to GST-BBS3 only in the presence of GTPγS (Fig. 3*F*). When the same procedure was performed on all combinations of GST-BBS3 and ARL13 variants, BBS3A73L but not BBS3T31R was bound to both ARL13G73L and ARL13T33N (Fig. 3*F*). As determined by size exclusion chromatography, BBS3A73L was bound to both ARL13G73L and ARL13T33N at a molecular ratio of 1:1 as the pulldown eluate cofractionated with a molecular weight of ~120 kDa (Fig. 3*G*). This confirmed that BBS3 only in a GTP-bound state interacts with ARL13 directly without relying on ARL13 nucleotide state (Fig. 3*F*). As determined by GEF assay, BBS3A73L but not BBS3T31R promoted ARL13 to bind to mant-GTP at a binding rate of 5.24× 10^-4^ (μmol mant-GTP/μmol ARL3/sec) (Fig. 3*H*). After the GEF effect of ARL13 on BBS3 was excluded, we conclude that BBS3^GTP^ activates ARL13^GDP^ by acting as an ARL13 GEF in cilia but not *vice versa* (Fig. 3*H*).

### A reservoir of the IFT-shed BBSome acts as an ARL13 effector in cilia

BBS3 undergoes GTPase cycle in cilia for shuttling between the ciliary membrane and matrix (49). In mammalian cilia, Rabl4/Ift27 activates Bbs3^GDP^, making Bbs3^GTP^ available at the ciliary tip for recruiting its GPCR-laden BBSome effector to load onto retrograde IFT trains for ciliary retrieval (34, 48). This does not hold true in *Chlamydomonas* cilia as loss of BBS3 in cilia causes PLD but not the BBSome to build up in cilia (49). As performed on cilia of *arl13-356; BBS3:YFP-TG, arl13-356; BBS3A73L:YFP-TG,* and *arl13-356; BBS3T31R:YFP-TG* cells, BBS3-YFP failed to immunoprecipitate the BBSome (represented by BBS1, BBS4, BBS5, BBS7, and BBS8) even in the presence of GTPγS and nor BBS3A73L-YFP did so, excluding the BBSome from being a BBS3 effector in cilia (Fig. 4*A*) (49). This notion was affirmed as the BBSome co-sedimented with none of BBS3A73L-YFP and the GTPγS- and GDP-loaded BBS3 in sucrose density gradients (Fig. 4*B*). When the assay was performed on cilia of *clip1; ARL13:HA:YFP-TG* and *clip1; ARL13G73L:HA:YFP-TG* cells, which lack BBS3 in cilia, GTPγS loading conferred ARL13-HA-YFP to recover the BBSome but not IFT-B1 (represented by IFT46 and IFT70) and so did for ARL13G73L-HA-YFP (Fig. 4*C*). None of the BBSome and IFT-B1 was recovered by the GDP-loaded ARL13-HA-YFP, identifying a reservoir of the BBSome, autonomous of IFT-B1 association, exists in cilia for ARL13^GTP^ to interact with (Fig. 4*C*). Supportively, ARL13G73L-HA-YFP resembled the GTPγS-but not GDP-loaded ARL13 in cofractionating with the BBSome in sucrose density gradients (Fig. 4*D*). According to the *in vitro* protein interaction assays, ARL13^GTP^ interacts with the BBSome via directly binding the BBSome subunit BBS1 as BBS1 was efficiently captured by ARL13G73L-HA-YFP but not ARL13T33N-HA-YFP (Fig. 4*E*). In sum, a reservoir of the IFT-shed BBSome acts as an ARL13 effector in cilia, and they bind each other in cilia in a BBS3-independent manner.

**Figure 4.**
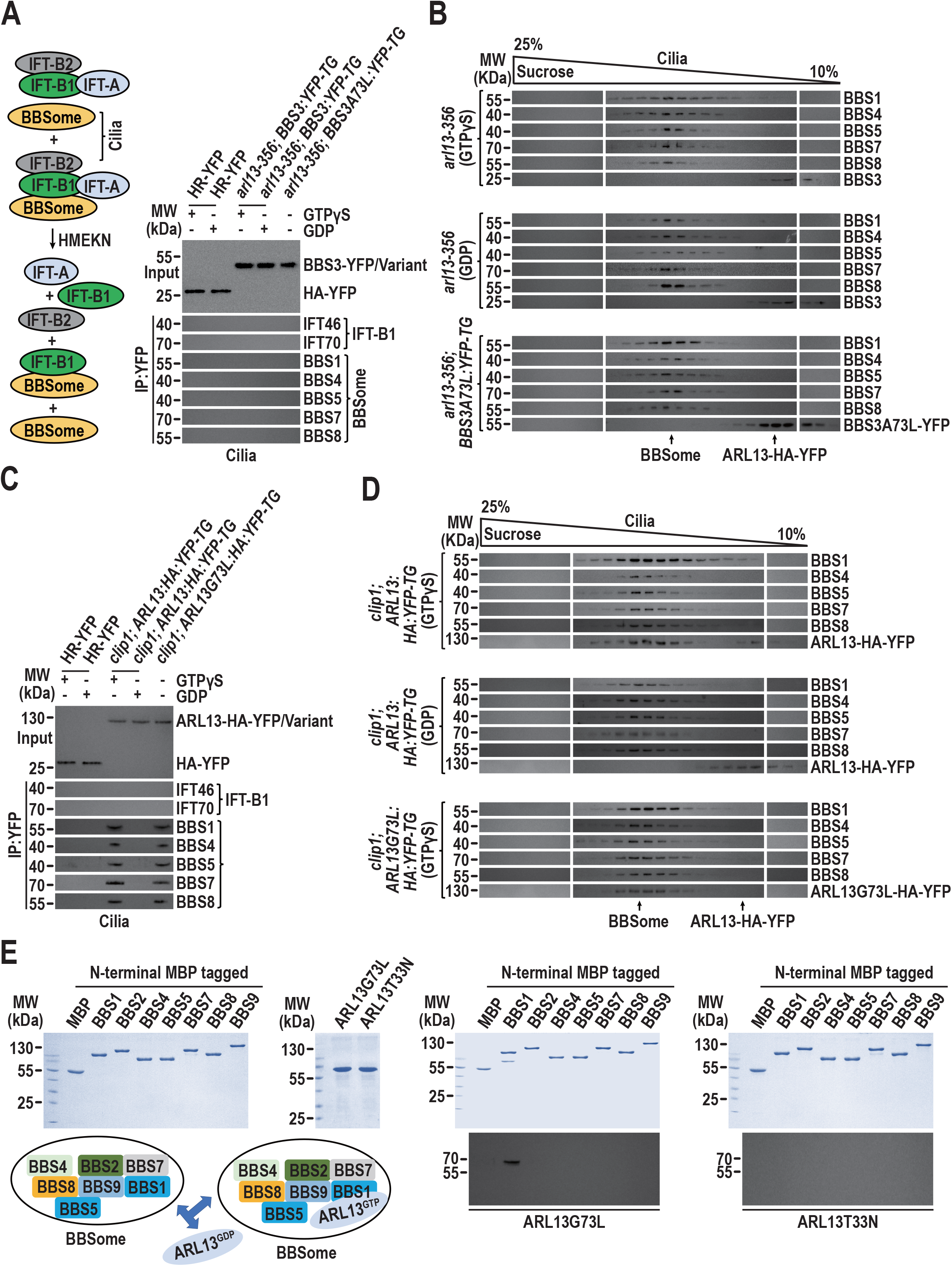
A reservoir of the IFT-shed BBSome acts as an ARL13 effector in cilia. (*A*). Schematic representation showing that IFT subcomplexes IFT-A, -B1, and -B2 remain to be separated from one another and only partial BBSome remains to bind to IFT-B1 in HMEKN buffer (Shown on left). Immunoblots of α-YFP-captured proteins from cilia of cells indicated on the top in the presence of GTPγS or GDP probed for the IFT-B1 subunits IFT46 and IFT70 and the BBSome subunits BBS1, BBS4, BBS5, BBS7, and BBS8 (Shown on right). Input was quantified with α-YFP by immunoblotting. (*B*). Immunoblots of sucrose density gradients of cilia of cells indicated on the left in the presence of GTPγS, GDP, or none of them probed with antibodies against the BBSome subunits BBS1, BBS4, BBS5, BBS7, and BBS8 and BBS3. (*C*). Immunoblots of α-YFP-captured proteins from cilia of cells indicated on the top in the presence of GTPγS or GDP probed for the IFT-B1 subunits IFT46 and IFT70 and the BBSome subunits BBS1, BBS4, BBS5, BBS7, and BBS8. Input was quantified with α-YFP by immunoblotting. (*D*). Immunoblots of sucrose density gradients of cilia of cells indicated on the left in the presence of GTPγS, GDP, or none of them probed with antibodies against the BBSome subunits BBS1, BBS4, BBS5, BBS7, and BBS8 and YFP. (*E*). Bacterially expressed MBP, MBP-BBS1, MBP-BBS2, MBP-BBS4, MBP-BBS5, MBP-BBS7, MBP-BBS8, and MBP-BBS9 (left) were mixed with ARL13G73L or ARL13T33N (second to the left) and complexes recovered on amylose beads were resolved by SDS-PAGE followed by Coomassie staining and immunoblotting with α-ARL13 (second to the right and right, respectively). A schematic representation of the direct interaction of ARL13^GTP^ with the BBS1 component of the BBSome was shown (lower left). For all panels, MW stands for molecular weight.

### ARL13^GTP^ recruits its IFT-shed BBSome effector to anchor to the ciliary membrane

As reflected by immunoprecipitation assays performed on cilia of *arl13; ARL13:HA:YFP-TG, arl13; ARL13Q73L:HA:YFP-TG,* and *arl13; ARL13T30N:HA:YFP-TG* cells, ARL13^GTP^ recovered BBS3^GTP^ and the BBSome simultaneously, while ARL13^GDP^ immunoprecipitated BBS3^GTP^ alone (Fig. 5*A*). BBS3 only in a GTP-bound form co-fractionated with ARL13 without relying on ARL13 nucleotide state; BBS3^GTP^, ARL13, and the BBSome co-sedimented only when ARL13 was GTP-bound, revealing that BBS3^GTP^ binding does not affect ARL13^GTP^ interaction with its BBSome effector in cilia (Fig. 5*B*). Interestingly, ARL13^GTP^ anchors to the ciliary membrane and binds its BBSome effector simultaneously, indicative of ARL13 to function as a molecular switch for recruiting the BBSome to anchor to the ciliary membrane. Reflecting this notion, ARL13 and the BBSome resided exclusively in the matrix of *clip1* cilia (Fig. 5*C*). This is easy to understand as, without its BBS3 GEF, ARL13 remains to be GDP-bound and fails to anchor to the ciliary membrane and nor binds the BBSome in the ciliary matrix (Fig. 2 and *SI Appendix,* Fig. S4*A* and *B*). In contrast, GTPγS loading conferred ARL13 to anchor to the ciliary membrane and, in this situation, partial BBSome became anchored to the ciliary membrane (Fig. 5*C* and *D*). Consistently, ARL13G73L-HA-YFP conferred partial BBSome to anchor to the ciliary membrane even when GDP was present to render the endogenous ARL13 GDP-bound in *clip1; ARL13G73L:HA:YFP-TG* cilia (Fig. 5*C*). Thus, upon activation, ARL13^GTP^ alone can bind and anchor its BBSome effector to the ciliary membrane. This notion was verified as ARL13G73L-HA-YFP and partial BBSome resided together with the GTPγS-loaded BBS3 in the membrane of *arl13-356; ARL13G73L:HA:YFP-TG* cilia (Fig. 5*D*). In the ciliary membrane, ARL13G73L-HA-YFP immunoprecipitated the GTPγS-loaded BBS3 alone in *arl13; bbs8; ARL13G73L:HA:YFP-TG* cells; the BBSome alone in *clip1; ARL13G73L:HA:YFP-TG* cells; and both the BBSome and the GTPγS-loaded BBS3 in *arl13-356; ARL13G73L:HA:YFP-TG* cells (Fig. 5*E*). Thus, upon anchoring to the ciliary membrane, ARL13^GTP^ remains to be bound to the BBSome in a BBS3^GTP^-independent manner (Fig. 5*F*).

**Figure 5.**
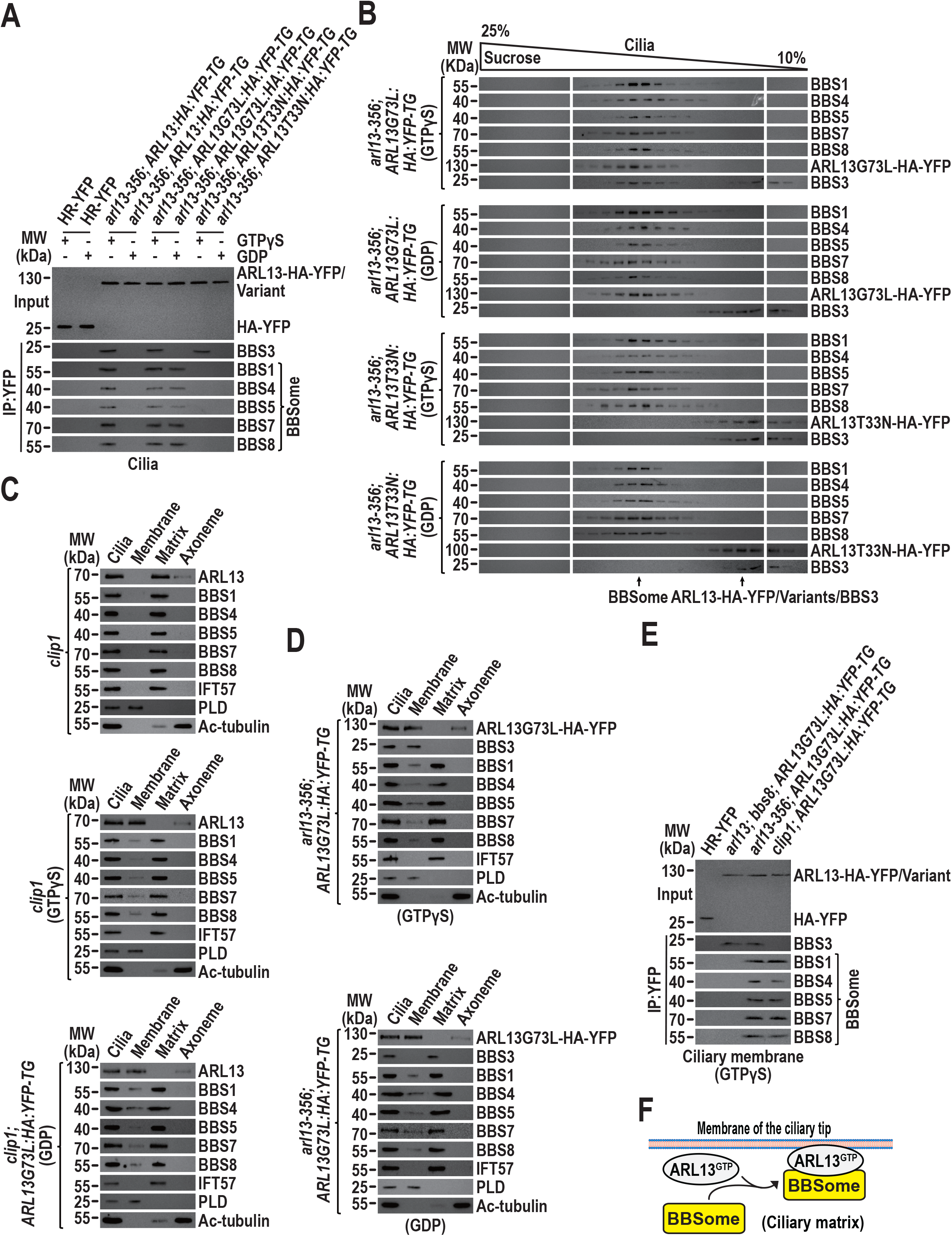
ARL13^GTP^ recruits its IFT-shed BBSome effector to anchor to the ciliary membrane. (*A*). Immunoblots of α-YFP-captured proteins from cilia of cells indicated on the top in the presence of GTPγS or GDP probed for BBS3 and the BBSome subunits BBS1, BBS4, BBS5, BBS7, and BBS8. Input was quantified with α-YFP by immunoblotting. MW: molecular weight. (*B*). Immunoblots of sucrose density gradients of cilia of cells indicated on the left in the presence of GTPγS or GDP probed with antibodies against the BBSome subunits BBS1, BBS4, BBS5, BBS7, and BBS8, BBS3, and YFP. MW: molecular weight. (*C*). Immunoblots of ciliary fractions of cells indicated on the left in the presence of GTPγS, GDP or none of them probed with α-ARL13, antibodies against the BBSome subunits BBS1, BBS4, BBS5, BBD7, and BBS8, α-IFT57 (ciliary matrix marker), α-PLD (ciliary membrane marker) and Ac-tubulin (axoneme marker). MW: molecular weight. (*D*). Immunoblots of ciliary fractions of cells indicated on the left in the presence of GTPγS or GDP probed with α-ARL13, α-BBS3, antibodies against the BBSome subunits BBS1, BBS4, BBS5, BBD7, and BBS8, α-IFT57 (ciliary matrix marker), α-PLD (ciliary membrane marker) and Ac-tubulin (axoneme marker). MW: molecular weight. (*E*). Immunoblots of α-YFP-captured proteins from the ciliary membrane of cells indicated on the top in the presence of GTPγS probed for BBS3 and the BBSome subunits BBS1, BBS4, BBS5, BBS7, and BBS8. Input was quantified with α-YFP by immunoblotting. MW: molecular weight. (*F*). Schematic presentation of how ARL13^GTP^ binds and anchors the BBSome to the ciliary membrane.

**Figure 6.**
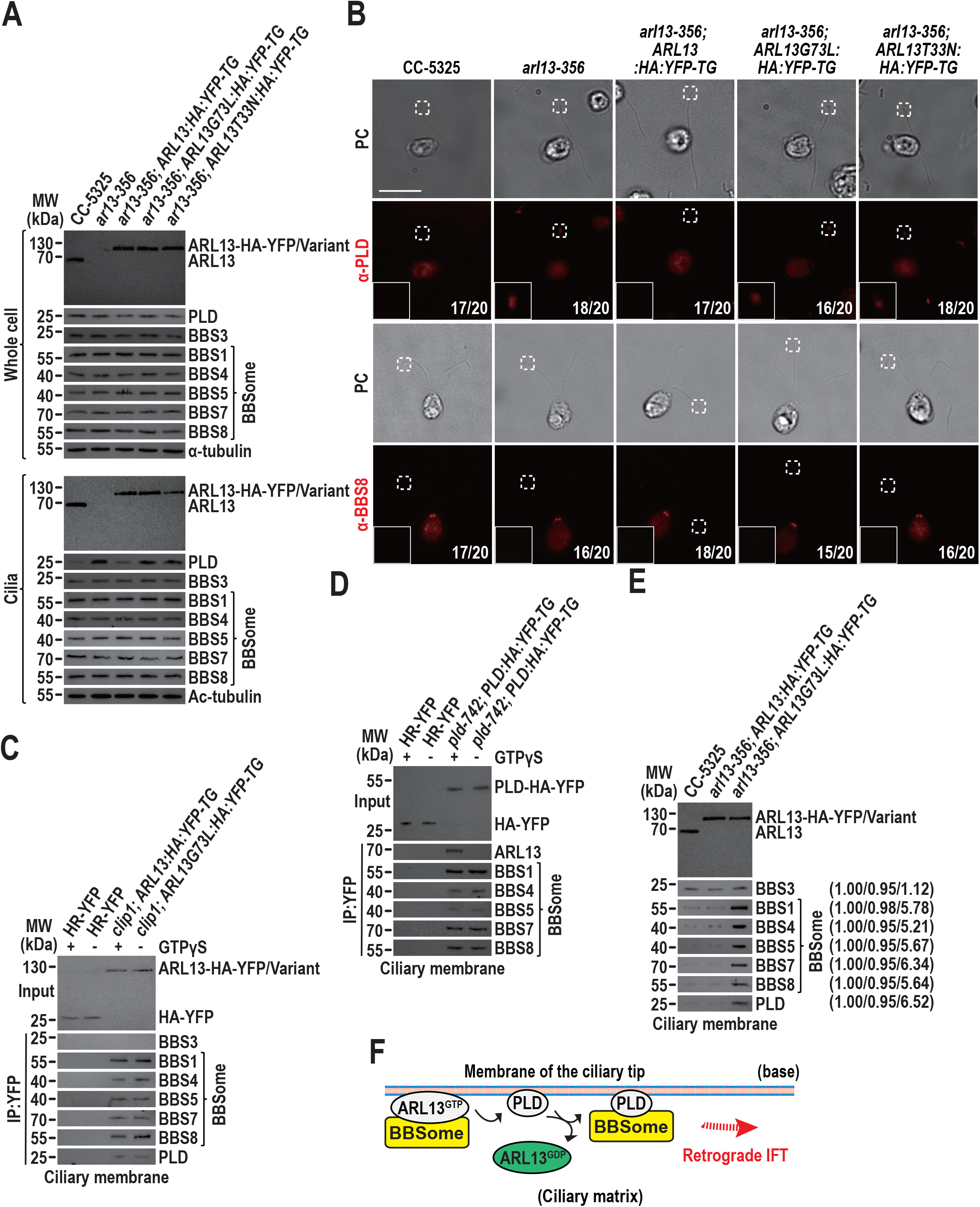
ARL13 GTP hydrolysis allows PLD-laden BBSome coupling with retrograde IFT at the ciliary tip. (*A*). Immunoblots of whole cell samples and cilia of cells indicated on the top probed with α-ARL13, α-BBS3, α-PLD, and antibodies against the BBSome subunits BBS1, BBS4, BBS5, BBS7, and BBS8. Alpha-tubulin and acetylated (Ac)-tubulin were used to adjust the loading of whole cell samples and cilia, respectively. MW: molecular weight. (*B*). Cells shown on the top stained with α-PLD (red) or α-BBS8 (red). PC images of cells were shown. Insets are 9 times enlargements of the ciliary tip indicated with a white box. Cell numbers out of 20 cells are listed for representing cells that accumulate PLD or BBS8 or not at the ciliary tip. Scale bar: 10 μm. (*C-D*). Immunoblots of α-YFP-captured proteins from the ciliary membrane of cells listed on the top in the presence of GTPγS or GDP probed for ARL13, BBS3, the BBSome subunits BBS1, BBS4, BBS5, BBS7, and BBS7, and PLD. Input was quantified with α-YFP by immunoblotting. MW: molecular weight. (*E*). Immunoblots of ciliary membrane of cells indicated on the top probed with α-ARL13, α-BBS3, the antibodies against the BBSome subunits BBS1, BBS4, BBS5, BBS7, and BBS8, and α-PLD. BBS3 was used as a loading control. The quantification of the band strengths normalized for those of BBS3 are shown in brackets on the right. The analysis is based on one experiment. MW: molecular weight. (*F*) Schematic presentation of how ARL13^GTP^ GTP hydrolysis releases PLD-laden BBSome to load onto retrograde IFT trains for migrating towards the ciliary base.

### ARL13 GTP hydrolysis allows PLD-laden BBSome coupling with retrograde IFT at the ciliary tip

ARL13 knockout does not alter cellular PLD abundance but resembled BBS3 ciliary loss in causing PLD accumulation mostly at the ciliary tip (Fig. 6*A* and *B*) (11, 49). PLD ciliary tip buildup is an outcome of the BBSome unavailability for PLD coupling in the ciliary membrane as BBSome ciliary content was normal without ARL13, consistent with the observation that the BBSome ciliary turnover remains normal in the absence of BBS3 in cilia (Fig. 6*A*) (11, 49). Notably, ARL13-HA-YFP but not its G73L and T33N versions restored PLD ciliary tip abundance back to normal (Fig. 6*A* and *B*). It is easy to understand as the ARL13T33N variant fails to anchor the BBSome to the ciliary membrane for PLD coupling (Fig. 5). In the case of the G73L variant that can anchor the BBSome to the ciliary membrane normally (Fig. 5), the PLD ciliary tip retention could be caused owing to the disability of the ciliary membrane-anchored BBSome to couple with PLD or the failure of the PLD-laden BBSome to load onto retrograde IFT trains for ciliary retrieval. The former assumption was invalid as ARL13G73L-HA-YFP and the GTPγS-loaded ARL13-HA-YFP immunoprecipitated PLD and the BBSome simultaneously in the ciliary membrane without relying on BBS3 (Fig. 6*C* and *SI Appendix,* Fig. S5*A*). To test the latter notion, we performed immunoprecipitation assays on the ciliary membrane of *pld-472; PLD:HA:YFP-TG* cells, which express PLD fused at its C-terminus to HA-YFP in the PLD-null CLiP mutant that we named *pld-472* (LMJ.RY0402.071742) (*SI Appendix,* Fig. S5*B*). PLD-HA-YFP recovered the BBSome and the GTPγS-but not GDP-loaded ARL13 (Fig. 6*D*). As compared to CC-5325 and *arl13-356; ARL13:HA:YFP-TG* cells, *arl13-356; ARL13G73L:HA:YFP-TG* cells contained approximately 6-fold higher PLD and the BBSome in the ciliary membrane, identifying GTP hydrolysis enables ARL13 to release the PLD-laden BBSome for retrograde IFT train loading (Fig. *6E* and *F*). The BBSome abundance remained normal even in cilia of *arl13-356; ARL13G73L:HA:YFP-TG* cells as determined by immunoblotting (Fig. 6*A*). Unlike PLD, it (represented by BBS8) was undetectable at the tip of *arl13-356; ARL13G73L:HA:YFP-TG* cilia by immunostaining (Fig. 6*B*). These may reflect the fact that only a very small partial of the total ciliary BBSome anchors to the membrane of the ciliary tip for PLD coupling (Fig. 5*C* and *D*).

## Discussion

The cilium is a sensing organelle of specialized membrane harboring transmembrane and membrane-tethered signaling proteins that act as IFT cargoes for exporting from and/or importing into cilia via IFT. To maintain their ciliary dynamics, they migrate from the ciliary tip towards base for ciliary retrieval via loading in a BBSome-dependent manner onto retrograde IFT trains. However, the molecular basis of the ciliary tip cargo-BBSome coupling process has remained uncertain. In this study, we found that ARL13 binds and anchors the post-remodeled BBSome as its effector to the ciliary membrane for PLD coupling at the ciliary tip (Fig. 7). During ARL13 GTPase cycle in cilia, BBS3 in a GTP-bound form activates ARL13 as an ARL13 GEF (Fig. 7). This BBS3-ARL13-mediated ciliary tip cargo-BBSome coupling mechanism closes a gap in our understanding of how cargo-BBSome coupling can occur at the ciliary tip.

**Figure 7.**
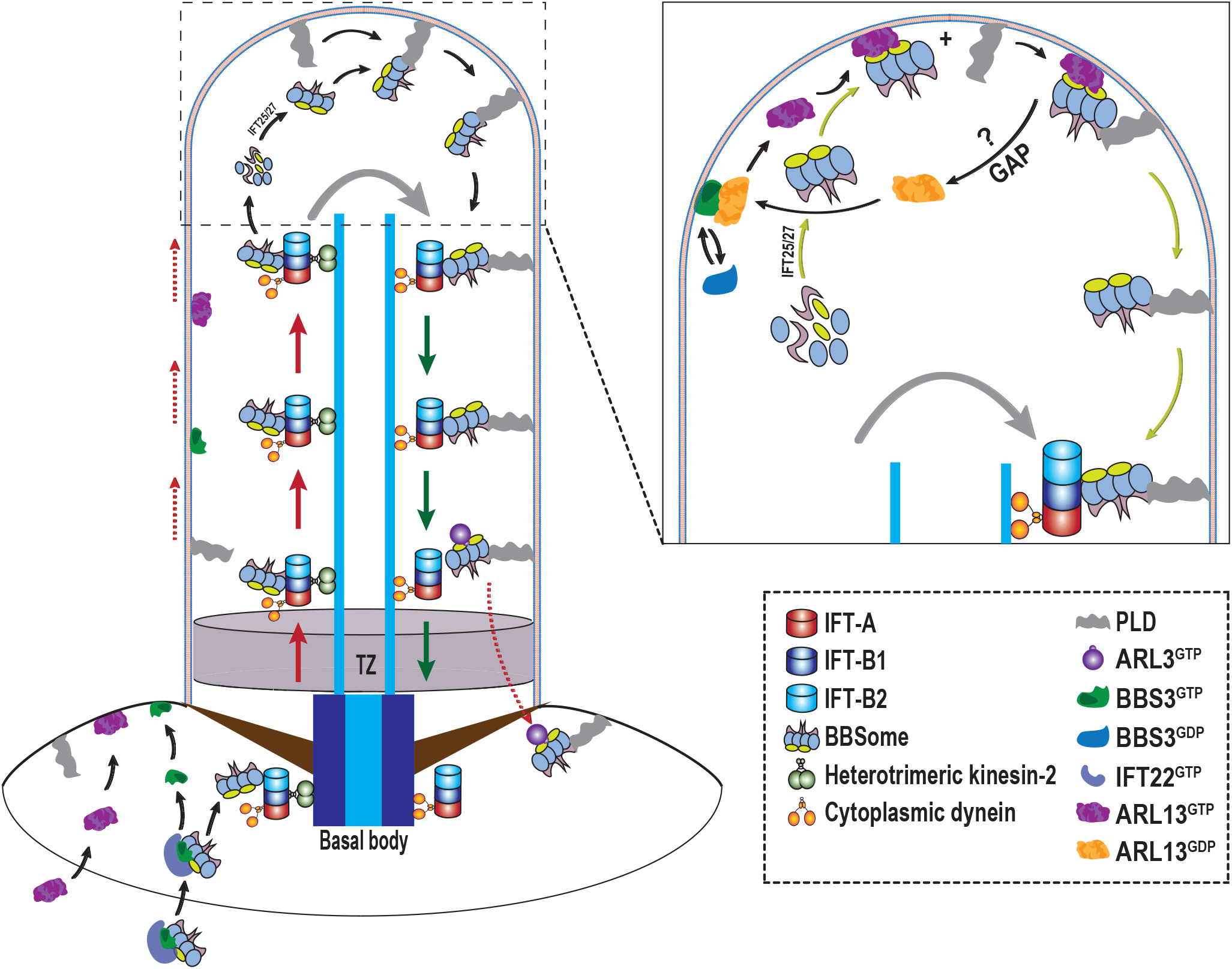
Hypothetical model for how BBS3 activates ARL13 to facilitate cargo-BBSome coupling on the membrane of the *Chlamydomonas* ciliary tip. BBS3 binds IFT22 to form the IFT22/BBS3 entity in cytoplasm (54). IFT22/BBS3 binds the BBSome via the interaction between BBS3 and the BBSome (54) Only when they both are GTP-bound, IFT22^GTP^/BBS3^GTP^ recruits the BBSome to the basal bodies (54). IFT22^GTP^/BBS3^GTP^/BBSome dissociates with one another at the basal bodies, allowing IFT22^GTP^ and the BBSome to integrate into anterograde IFT trains and BBS3^GTP^ to anchor to the membrane for diffusing into cilia followed by GTPase cycle (28, 54). ARL13 in a GTP-bound state targets to the basal bodies at physiological conditions. At the basal bodies, ARL13^GTP^ anchors to the membrane for diffusing into cilia followed by GTPase cycle. GTPase cycle confers them to shuttle between the ciliary membrane (BBS3^GTP^ and ARL13^GTP^) and matrix (BBS3^GDP^ and ARL13^GDP^) dynamically. At the ciliary tip, BBS3^GTP^ activates ARL13 as an ARL13 GEF. ARL13^GTP^ binds and recruits the post-remodeled BBSome as an ARL13 effector to anchor to the ciliary membrane. This spatiotemporally enables the BBSome to couple with the ciliary membrane-tethered PLD. Thereafter, GTP hydrolysis of ARL13^GTP^ allows ARL13 to dissociate with the PLD-laden BBSome, releasing it to load onto retrograde IFT trains for migrating towards to the ciliary base. Upon migrating to the proximal ciliary region right above the TZ, the PLD-laden BBSome dissociates with retrograde IFT trains and moves across the TZ for ciliary retrieval via the ARL3-mediated outward BBSome TZ diffusion pathway (28). During ARL13 GTPase cycle, a ciliary ARL13 GAP is potentially required (?). IFT25/27 promotes BBSome reassembly during its remodeling at the ciliary tip (41).

### ARL13 uses the BBSome as its effector at the ciliary tip

As compared to other species, *C. reinhardtii* facilitates an unbiased biochemical analysis of isolated cilia and cell bodies (50, 51). By taking advantage of this superiority, we found that IFT22/BBS3 binds the BBSome via a direct interaction between BBS3 and the BBSome and recruits the BBSome to the basal bodies only when they both are GTP-bound, mediating BBSome ciliary dynamics by controlling its basal body amount available for entering cilia (54). Upon translocating to the basal bodies from cytoplasm, BBS3^GTP^ dissociates with IFT22^GTP^ for anchoring to the membrane for diffusing into cilia (49, 54). Inside cilia, BBS3^GTP^ undergoes GTPase cycle but itself does not use the BBSome as an effector (Fig. 4*A* and *B*) (49). It has been known that ARL3 uses the BBSome as its effector when they both position to the proximal ciliary region right above the TZ (28). At this ciliary region, ARL3^GTP^ binds and recruits the PLD-laden BBSome, autonomous of retrograde IFT, to diffuse through the TZ for ciliary retrieval (28). In this study, the BBSome was identified to function as an ARL13 effector in cilia (Fig. 4*C-E*), while this GTPase/effector process is restricted to happen at the ciliary tip (Fig. 6*A* and *B*). ARL3 and ARL13 fail to bind the BBSome in cytoplasm probably owing to the IFT22/BBS3 binding to the BBSome, which occupies the BBSome binding site(s) located in the BBS1 subunit for ARL3 and ARL13 (Fig. *4E*) (52, 54). In rodent cells, the BBSome was implicated to be a BBS3 effector based on biochemical assays performed on whole cell samples (48). Our current finding does not necessarily disapprove the BBSome to be a BBS3 effector in rodent cilia, while *Chlamydomonas* cells direct BBSome trafficking between different cellular compartments and even at different locations of the same organelle as the cilium via distinct GTPase pathways.

### BBS3 activates ARL13 as an ARL13 GEF to anchor the BBSome to the membrane of the ciliary tip

The BBSome undergoes IFT for maintaining its ciliary dynamics. During this process, it detaches from anterograde IFT for remodeling at the ciliary tip followed by loading onto retrograde IFT for migrating towards the ciliary base (41). Thus, cargo-BBSome coupling at the ciliary tip is critical as many ciliary transmembrane and membrane-tethered signaling molecules i.e. GPCRs and PLD rely on the post-remodeled BBSome for proper ciliary retrieval via retrograde IFT (34, 49). We previously found that *Chlamydomonas* BBS3 facilitates PLD coupling with the BBSome at the ciliary tip and ARL13 resembled BBS3 in facilitating PLD coupling with the BBSome in cilia (11, 49). In this study, we found that ARL13 in a GTP-bound state anchor to the membrane for diffusing into cilia (Fig. 1*A* and *B*), resides along the whole length of cilia by tethering to the ciliary membrane (Fig. 1*D* and *E*), undergoes GTPase cycle with BBS3 acting as an ARL13 GEF (Fig. 1 and 3), and recruits the post-remodeled and ciliary matrix-residing BBSome as its effector to anchor to the ciliary membrane (Fig. 4 and 5). This enables the BBSome to become spatially adjacent to its cargo PLD on the membrane of the ciliary tip, facilitating PLD coupling with the BBSome (Fig. 6). The *Chlamydomonas* cell has developed such an elaborated system to restrict ARL13 to bind the BBSome in the matrix of the ciliary tip where the BBSome drops off anterograde IFT for remodeling (41). ARL13 mediates ciliary membrane anchoring of the BBSome as its effector at the ciliary tip for transmembrane and membrane-tethered signaling proteins to couple with, playing a critical role in maintaining proper ciliary signaling (Fig. 7). It is also worth to note that ARL13^GDP^ conversion to ARL13^GTP^ may occur with the aid of an unknown ARL13 GAP in cilia as ciliary extracts dramatically elevate ARL13 GTPase activity (Fig. 1*G* and 7).

### BBS3 GTPase cycle in cilia

BBS3 enters cilia as being a GTP-bound state and undergoes GTPase cycle (48, 49). *Chlamydomonas* BBS3 differs from its mammalian orthologues in that it contains an A73 in the G1 domain rather than a Q73 essential for GTP hydrolysis of small GTPases (49). Interestingly, BBS3 resembles BBS3A73Q in hydrolyzing GTP with low intrinsic GTPase activity (*SI Appendix,* Fig. S6*A*). There is no clue thus far for how BBS3 GTP hydrolysis could initiate without Q73, while A73 unlikely influence BBS3 GTPase activity. Since ciliary extracts failed to elevate BBS3 GTPase activity (*SI Appendix,* Fig. S6*A*), BBS3^GTP^ conversion to BBS3^GDP^ is assumed to be very slow as a BBS3 GAP may not exist in *Chlamydomonas* cilia. The low intrinsic GTPase activity of BBS3 reveals that BBS3 could mainly exist as a GTP-bound form in cilia, consistent with our previous observation (49). IFT27 binds its partner protein IFT25 to exist as a heterodimeric IFT25/27 entity cross-species (56, 57). Inconsistent with the rodent Ift25/27 that was implicated to provide a docking site for the BBSome to link to IFT machinery (42) or act as a GEF for BBS3 (34), *Chlamydomonas* IFT25/27 in either of IFT27 nucleotide state fails to activate BBS3 *in vitro (SI Appendix,* Fig. S6*B* and *C*) and its role in promoting BBSome reassembly during BBSome remodeling at the ciliary tip has been proven (41). This raised an interesting question, namely which factor other than IFT25/27, if desirable, contributes to activate BBS3 in cilia of *C. reinhardtii.* We currently hold no answer for this question, while the BBS3 GEF in *Chlamydomonas* cilia deserves to be identified in the future.

### Implications for BBS molecular basis

Our biochemical data identify only a very small partial of ciliary BBSomes jump off anterograde IFT trains at the ciliary tip (Fig. 5*C* and *D*). For these unshed from IFT trains, they are supposed to remain to be bound to IFT for moving along the axoneme bidirectionally. This reservoir of IFT-shed BBSome undergoes remodeling process and anchors to the ciliary membrane for signaling protein cargo coupling (Fig. 7). In mammals, GPCRs in Hh signaling pathway move out of cilia via the IFT/BBS system (35, 48). Uncoupling of GPCRs with the BBSome could occur at the ciliary tip if the BBSome fails to anchor to the membrane of the ciliary tip due to certain mutations in *BBS3, ARL13b, IFT25,* and *IFT27* genes (34, 42, 43). Thus, unraveling this intricate ciliary tip cargo-BBSome coupling mechanism will allow us to better understand the molecular basis of BBS. If this holds true in humans, why ARL13b-related JBTS shares overlapping phenotypes with BBS disorders in humans can be satisfactorily explained.

## Materials and methods

### Antibodies, Chlamydomonas strains, and culture conditions

Antibodies including α-BBS1, α-BBS3, α-BBS4, α-BBS5, α-BBS7, α-BBS8, α-IFT22, α-IFT38, α-IFT43, α-IFT46, α-IFT57, α-IFT70, α-IFT139, and α-PLD have been described previously and were listed in *SI Appendix, Table S1* (41, 49, 54, 58). Rabbit-originated antibodies against ARL13 were generated with bacterially expressed ARL13 of full amino acid length by Beijing Protein Innovation, LLC (Beijing). Antibodies recognizing YFP (mAbs 7.1 and 13.1, Roche), α-tubulin (mAb B512, Sigma), and acetylated-tubulin (mAb 6-11B-1, Sigma-Aldrich) were commercially bought (*SI Appendix, Table S1).* The strain CC-5325 and the *arl13-356* (LMJ.RY0402.181356), *clip1* (LMJ.RY0402.149010), and *pld-742* (LMJ.RY0402.071742) mutants were bought from the *Chlamydomonas* Library Project (CLiP; https://www.chlamylibrary.org/allMutants) (59). The BBS8-null mutant *bbs8* has been reported previously (55). All the strains used in this study are listed in *SI Appendix, Table S2*.

Various experimental protocols were applied in this study, and most of the experiments were briefly introduced in the text to make the content easier to be understood. The details of each protocol are available in *SI Appendix*.

### Statistical analysis

Statistical analysis was done with GraphPad Prism 8.30 (GraphPad Software). For *in vitro* GEF and small GTPase assays, the data were presented as mean ± S.D. from three duplicates in one experiment.

## Data availability

Data supporting the findings of this paper are contained within the paper and SI Appendix.

## Acknowledgments

We thank Drs. Karl Lethtreck and Jin Dai at the University of Georgia for constructive discussions about the role of ARL13 in cargo-BBSome coupling. Research reported in this publication was supported by National Natural Science Foundation of China Grant 32070698 to Z-C.F. and National Natural Science Foundation of China Grant 32200558 and China Postdoctoral Science Foundation Grant 2021M702457 to Y-X.L. The founders have no role in study design, data collection and analysis, decision to publish, or preparation of the manuscript.

## Author contributions

Z-C.F. designed research; Y-X.L., W-J.L., R-K.Z. and S-N.S. performed research; Z-C.F. and Y-X.L. analyzed data; and Z-C.F. wrote the paper.

