## Supplementary materials for "Unraveling the Intricate Cargo-BBSome Coupling Mechanism at the Ciliary Tip"

Zhen-Chuan Fan

**This PDF file includes:**

Materials and methods
Figures S1 to S6
Tables S1 to S3
Legends for Movies S1 to S3
SI References

**Other supplementary materials for this manuscript include the following:**

Movies S1 to S3

### Materials and methods

#### Plasmids and transgenic strain generation

Expression vectors were constructed on pBKS-gARL3-HA-YFP-Ble that contained a HA-YFP coding sequence followed immediately downstream by a sequence encoding the Rubisco 3'-UTR and the bleomycin (*Ble*, zeocine resistant gene) cassette (1). Genomic DNA of *Chlamydomonas* cells was extracted and purified using a Wizard® Genomic DNA Purification Kit (Promega, Beijing) following the kit's protocol. The vectors expressing BBS3-YFP, BBS3A73L-YFP, and BBS3T31R-YFP have been reported previously (2, 3). To generate the ARL13-HA-YFP expression vector, a 3,919-bp ARL13 DNA fragment containing 1,000-bp promoter sequence was amplified from genomic DNA by using the primer pair gARL13-FOR and gARL13-REV as listed in Table S3. This fragment was inserted into pBluescript II KS(+) vector linearized by *Xba*I and *Eco*RI, resulting in pBKS-gARL13. To obtain ARL13ΔN15, two DNA fragments were amplified by using primer pairs gARL13-FOR and gARL13ΔN15-REV and gARL13ΔN15-FOR and gARL13-REV as listed in Table S3 and inserted into the pBluescript II KS(+) vector linearized by *Xba*I and *Eco*RI, resulting in pBKS-gARL13ΔN15. Site-directed mutagenesis was performed to introduce the G73L and T33N mutations into pBKS-gARL13 or pBKS-gARL13ΔN15 by using the primer pairs ARL13G73L-FOR and ARL13G73L-REV and ARL13T33N-FOR and ARL13T33N-REV, respectively, as listed in Table S3. These *Xba*I and *Eco*RI double digested ARL13/variant DNA fragments and the HA-YFP-Ble DNA fragment cut from pBKS-gARL3-HA-YFP-Ble by *Eco*RI and *Kpn*I were inserted into pBluescript II KS(+) vector linearized by *Xba*I and *Kpn*I by three-way ligation, resulting in pBKS-gARL13-HA-YFP-Ble, pBKS-gARL13G73L-HA-YFP-Ble, pBKS-gARL13T33N-HA-YFP-Ble, pBKS-gARL13ΔN15-HA-YFP-Ble, pBKS-gARL13ΔN15G73L-HA-YFP-Ble, pBKS-gARL13ΔN15T33N-HA-YFP-Ble, respectively. To clone PLD-HA-YFP, a 2,956-bp PLD DNA fragment containing 1,000-bp promoter sequence was amplified by using the primer pair gPLD-FOR and gPLD-REV as listed in Table S3. Next, the *Xba*I and *Eco*RI double digested PLD fragment and the HA-YFP-Ble fragment cut from pBKS-gARL3-HA-YFP-Ble by *Eco*RI and *Kpn*I were inserted into pBluescript II KS(+) vector linearized by *Xba*I and *Kpn*I by three-way ligation, resulting in pBKS-gPLD-HA-YFP-Ble. All newly obtained constructs were affirmed by direct nucleotide sequencing. BBS3 miRNA vector pMi-ARL3 has been described previously (2). The screening of BBS3 miRNA cells was initiated by checking the cellular level of the target proteins through immunoblotting of whole cell extracts with BBS3 antibody. The miRNA strains showing a reduced level of the target proteins were selected for further phenotypic analysis. *C. reinhardtii* was transformed with the constructed vectors as shown above by the electroporation method as described previously (4) and the transformants were selected on TAP plates with 20 µg/ml paromomycin (Sigma-Aldrich), 15 µg/ml bleomycin (Invitrogen) or both antibiotics with 10 µg/ml paromomycin and 5 µg/ml bleomycin. The positive transformants were identified and the target proteins were quantified directly by immunoblotting as described below. The screening of BBS3 miRNA cells was initiated by checking the cellular level of the target protein through immunoblotting of whole cell extracts with the BBS3 antibody. The miRNA strains showing a reduced level of the target protein were selected for further phenotypic analysis.

#### ***Chlamydomonas* strain crossing**

The ARL13- and BBS8-double null mutant *arl13; bbs8* and the *clip1; bbs8* mutant were generated by crossing the relevant single mutants as described previously (1). In briefly, 20 ml of each of *arl13-356*, *clip1*, and *bbs8* mutants were grown in M1 medium to a density of $2\times10^6$ cells/ml, washed with M1-N (without nitrogen) medium for three times, transferred to 20 ml of M-N medium followed by aeration overnight (~12 hrs) under constant light at room temperature. Gametes of two opposite mating types were mixed in flask and incubated under light for 2 hrs. The cells siting on the bottom of the flask were transferred to M1 plate containing 4% agar, air-dried, and incubated overnight under continuous light. Cells were then incubated in the dark for a week prior to being moved to incubate at -20°C for two days. Afterward, cells were moved to incubate under continuous light at room temperature for at least ten days. The *arl13; bbs8* double mutant was screened by detecting the loss of ARL13 and BBS8 proteins via immunoblotting of whole cell samples with both ARL13 and BBS8 antibodies. The *clip1; bbs8* mutant was screened by the same way but via immunoblotting of ciliary extracts with both BBS3 and BBS8 antibodies.

#### **Isolation of cilia and cell bodies**

Isolation of *Chlamydomonas* cilia and cell bodies was done according to our protocol reported previously (5). If not otherwise specified, *Chlamydomonas* cells were grown in 10 liters of TAP medium until they reach a density of $10^8$ cells/ml. The cells were then collected by centrifugation at 1000 ×*g* for 15 min at room temperature, resuspended in 150 ml TAP medium (pH7.4), and incubated for 2 hrs under strong light with bubbling. Afterward, the cells were added with 0.5 M acetic acid for adjusting the pH value to 4.5 for deciliating the cells within 30-60 sec. Next, 0.5 M KOH was added to adjust the pH value back to 7.4. *Chlamydomonas* cell bodies were collected as pellets by centrifugation at 600 ×*g* 4°C for 5 min. The supernatant was centrifuged again at 12,000 ×g 4°C for 10 min and the pellets were collected as cilia. Cilia were repeatedly washed with HMDEKN buffer (30 mM Hepes [pH 7.4], 5 mM $MgSO_4$, 1 mM DTT, 0.5 mM EGTA, 25 mM KCl, 125 mM NaCl) supplemented with protein inhibitors (PI) (1 mM PMSF, 50 μg/ml soy-bean trypsin inhibitor, 1 μg/ml pepstatin A, 2 μg/ml aprotinin, and 1 μg/ml leupeptin) by centrifugation at 12,000 ×*g* 4°C for 10 min until the green color disappeared completely.

#### **Preparation of ciliary fractions**

Ciliary membrane, matrix, and axoneme were prepared according to our protocol described previously (2). Briefly, cilia suspended in HMDEKN buffer plus PI underwent three cycles of frozen-

and-thaw in liquid nitrogen. After centrifugation at 12,000 ×g at 4°C for 15 min, the supernatant was collected as the ciliary matrix. The pellets composed of ciliary membrane and axoneme were resuspended in HMEDKN buffer plus PI supplemented with 1% nonidet P-40 (NP-40) followed by sitting on ice for 30 min. The supernatant and pellets were collected as ciliary membrane and axonemal, respectively, by centrifugation at 12,000 ×g at 4°C for 10 min.

**Sucrose density gradient centrifugation assay**

Sucrose density gradient centrifugation of cell body and ciliary extracts were done according to our protocol reported previously (6). In brief, linear 12 ml of 10-25% sucrose density gradients were prepared in HMDEKN. Cilia and the cell bodies suspended in HMDEKN plus PI supplemented with 1% NP-40 were frozen-and-thawed for three cycles in liquid nitrogen. After centrifugation at 12,000 ×*g* at 4℃ for 10 min, the non-soluble debris was removed and 700 μl of supernatants were loaded on the top of the gradients followed by centrifugation at 38,000 rpm at 4℃ for 14 hrs in a SW41Ti rotor (Beckman Coulter). Afterward, the gradients were collected as 24 fractions of 0.5 ml aliquots. To detect protein co-sedimentation pattern, 20 μl of each fraction was loaded into 12% SDS-PAGE gel for electrophoresis and analyzed by immunoblotting as described below. The standards used to calculate S-values were BSA (4.4S), aldolase (7.35S), catalase (11.3S), and thyroglobulin (19.4S). If needed, 20 mM GTPγS or GDP were present.

**Immunoblotting**

Immunoblotting was performed on whole cell, cell body, and ciliary samples and the immunoprecipitated protein samples according to our protocol reported previously (7). Samples were mixed with Laemmli SDS sample buffer and boiled for 5 min followed by centrifugation at 2,500 ×*g* for 5 min to remove the debris. If not otherwise specified, either 20 µg (whole cell and cell body samples and the immunoprecipitated protein samples) or 100 µg (ciliary samples) of total protein from each sample was loaded in the 12% SDS-PAGE gel for electrophoresis before being transferred to nitrocellulose (NC) membrane. The NC membranes were then blocked in 5% non-fat dry milk in TBS (10 mM Tris, pH 7.5, 166 mM NaCl) plus 0.05% Tween-20 and incubated with primary antibodies diluted in the blocking solution for 2 hrs at room temperature. Following washing for three times with TBS plus 0.05% Tween-20, the membranes were incubated with HRP-conjugated secondary antibodies for 2hrs at room temperature. Following the same washing for five times, chemiluminescence were used to detect the primary antibodies by using the Mini Chemiluminescent/Fluorescent Imaging and Analysis System (Sagecreation). Primary and secondary antibodies were diluted for immunoblotting with a ratio as shown in Table S1. If needed, ImageJ software (version 1.42g, National Institutes of Health) was applied for quantifying the target proteins by measuring the immunoblot intensity according to our protocol reported previously (3). The immunoblot intensity was normalized to the intensity of a loading control protein.

**Immunoprecipitation**

Immunoprecipitation was performed on the cell body, cilia, the ciliary membrane, and the ciliary matrix, respectively, according to our protocol reported previously (2). Briefly, samples prepared from *Chlamydomonas* cells expressing HA-YFP, YFP-tagged BBS3 and its variants, HA-YFP-tagged ARL13 and its variants, and HA-YFP-tagged PLD were resuspended in HMDEKN or DTT-deprived HMEKN buffer plus PI supplemented with 1% NP-40 and frozen and thawed for three cycles in liquid nitrogen. After centrifugation at 70,000 ×*g* at 4°C for 1 hr, the supernatants were collected for agitating with 5% BSA-pretreated camel anti-YFP antibody-conjugated agarose beads (V-nanoab Biotechnology) for 2 hrs at 4°C. By centrifugation at 2,500 ×*g* at 4°C for 2 min, the beads were washed with 1 ml HMDEKN or HMDEKN buffer for three times, with 1ml HMDEK or HMEK buffer containing 50 ml NaCl twice, and with 1ml HMDEKN or HMEKN buffer twice. The beads were then added with 100 μl of 0.1 M Glycine buffer (pH2.5) for incubation for 2 min at room temperature followed by centrifugation at 2,500 ×*g* at 4°C for 2 min for collecting the supernatant. The supernatants were then mixed with Laemmli SDS sample buffer and boiled for 5 min before centrifugation at 2,500 ×*g* for 5 min. The immunoprecipitants in the supernatants were collected for immunoblotting. If needed, 20 mM GTPγS or GDP were present.

**Liposome flotation assays**

Total RNA of *Chlamydomonas* cells was extracted and purified using Eastep® Super total RNA Extract Kit (Promega, Shanghai). Five micrograms of RNA were reverse transcribed at 42 °C for 1 h using M-MLV Reverse Transcriptase (Promega) and oligo(T)18 primers (Takara). ARL13 and ARL13ΔN15 cDNAs were amplified by PCR using primer pairs cARL13-FOR and cARL13-REV and cARL13ΔN15-FOR and cARL13-REV, respectively, as listed in Table S3. The PCR reactions were performed at 95 °C for 5 min followed by 30 cycles of 95°C for 20 sec, 61°C for 20 sec, and 72°C for 4 min. The DNA fragments obtained were inserted into the *Eco*RI and *Xho*I sites of pET-28a (Novagen) for generating pET-28a-cARL13 and pET-28a-cARL13ΔN15. G73L and T33N mutations were introduced into ARL13 and ARL13ΔN15 cDNAs by site-directed mutagenesis using primer pairs ARL13G73L-FOR and ARL13G73L-REV and ARL13T33N-FOR and ARL13T33N-REV, respectively, as listed in Table S3, resulting in pET-28a-cARL13G73L, pET-28a-cARL13T33N, pET-28a-cARL13ΔN15G73L, and pET-28a-cARL13ΔN15T33N. These plasmids were transformed into *Escherichia coli* BL21(DE3) and the C-terminal 6×His tagged ARL13 or its variants were purified with Ni Sepharose$^{TM}$ 6 Fast Flow beads (GE Healthcare) and, thereafter, the His-tag was removed by thrombin (Solarbio, Beijing) at the concentration of 3 U/mg as described in our previous report (3). Liposome flotation assays were performed according to our previous report (1). In brief, rehydrated lipid films in the lipid reconstitution buffer (30 mM Tris, 150 mM NaCl, 2 mM $MgCl_2$, 2 mM DTT, pH 7.5) were mixed by centrifuging at 1,000 rpm for 1 hr at room temperature. Liposomes were generated from re-suspended lipids with an extruder (200 nm pore size, 20 strokes manually, Avanti). Thereafter, 70 µg of purified ARL13 (100 µM) was incubated in the presence of 500 µM GTPγS or GDP. $MgCl_2$ was added to samples to reach a final concentration of 100 mM and the ARL13 protein was separated from the excess of nucleotides by Zeba Spin desalting columns (Thermal Scientific). Afterward, 75 µl of liposomes and lipid reconstitution buffer were mixed to reach a final volume of 150 µl and incubated for 2 hrs at room temperature. The reactions were added with 100 µl 75% sucrose solution, 200 µl 25% sucrose, and 50 µl lipid reconstitution buffer, and centrifuged at 240,000 ×g for 1 hr (Beckman TLS-55 rotor, OptiMax MAX-XP benchtop centrifuge). After the centrifugation, 50 µl solutions were collected from the top of the centrifuge tube and subjected to immunoblotting analysis with α-ARL13. The purified ARL13 variants were used for liposome flotation assays without nucleotide preloading. For immunoblotting, 1 µg of ARL13/variant protein was loaded for evaluating their binding/input ratio shown as percentile.

**Fixed imaging**

Immunofluorescence staining was conducted according to our previous report (8). Briefly, after *Chlamydomonas* cells growing in M1 media were washed once with 5 mM EGTA, they were seeded to 0.1% polyethyleneimine-coated coverslips for 8 min. Cells were permeabilized and fixed with ice-cold methanol twice each for 10 min and rehydrated with phosphate buffered saline (PBS). Thereafter, cells were blocked in blocking buffer (5% BSA, 1% cold water fish gelatin, and 10% goat serum in PBS) for 4 hrs and incubated with primary antibodies in blocking buffer for 2 hrs. Cells were then washed ten times in PBS followed by incubating with secondary antibodies in blocking buffer for 1 hr. After washing additional ten times in PBS, the coverslips were mounted with SlowFade Antifade reagent (Molecular Probes) and images were captured with an Olympus IX83 inverted fluorescent microscopy equipped with a back illuminated scientific CMOS camera (Prime 95B, Photometrics), a 100×/1.40 NA oil objective lens (Olympus), and 561-nm laser from Coherent OBIS Laser Module. All images were acquired and processed with CellSens Dimension (version 2.1, Olympus). The primary antibodies against ARL13, BBS3, PLD, and BBS8 and the secondary antibody Alexa-Fluor594-conjugated goat anti-rabbit (Molecular Probes) were listed in Table S1 with their suggested dilutions. The staining was performed at room temperature.

**Total internal reflection fluorescence (TIRF) microscopy and kymogram analysis**

Cells expressing YFP-tagged ARL13 and its variants (10 µl) growing in M1 medium were placed onto the larger coverslip (24×60-mm No.1.5) followed by settling for 2 min at room temperature. A ring of petroleum jelly was then added around the droplet and a smaller coverslip (22×22-mm No.1.5) containing 5 µl of 10 mM HEPES, pH 7.4 and 5 mM EGTA was inverted onto the larger one to generate a sealed observation chamber. TIRF microscopy was applied to visualize the motility of YFP-tagged ARL13 and its variants in cilia. Videos were captured with TIRF (15 Hz) for

~20 s by using the software CellSens Dimension (version 2.1, Olympus) at 15 frames per second (fps). For capturing video, an Olympus IX83 inverted fluorescent microscopy equipped with a through-the-objective TIRF system, a 100×/1.49 NA TIRF oil immersion objective len (Olympus), a back illuminated scientific CMOS camera (Prime 95B, Photometrics), and 488-nm laser from Coherent OBIS Laser Module was used according to our previous report (3). The videos were processed with CellSens Dimension (version 2.1, Olympus) for generating kymographs according to our protocol reported previously (7).

**Protein-protein interaction assay**

Bacterially expressed N-terminal MBP-tagged BBS1, BBS2, BBS4, BBS5, BBS7, BBS8, and BBS9 proteins were purified as described previously (1). To determine ARL13 interaction to the BBSome, 100 µg of MBP and the MBP-tagged BBS proteins were individually mixed with 100 µg of bacterially expressed ARL13G73L or ARL13T33N (see above) to form a combination of 16 reactions. After incubated for 2 hrs at room temperature, mixtures were purified with Dextrin Sepharose™ High Performance MBP-tagged protein purification resin (GE Healthcare). Ten micrograms of proteins from elutes was loaded on 12% SDS-PAGE gels for electrogenesis. Gels were then visualized with Coomassie blue staining. Immunoblotting assay was also performed to verify the interaction between BBS proteins and ARL13 variants with α-ARL13. The vectors expressing GST, GST-BBS3, GST-BBS3A73L, and GST-BBS3T31R have been reported previously (3). The A73Q mutation was introduced into pGEX-2T-cBBS3 by site-directed mutagenesis using primer pairs BBS3A73Q-FOR and BBS3A73Q-REV as listed in Table S3, resulting in pGEX-2T-cBBS3A73Q. These plasmids were transformed into *Escherichia coli* BL21(DE3) and the N-terminal GST tagged BBS3 or its variants were purified with GST SpinTrap™ Purification Module (GE Healthcare) as described in our previous report (3). 100 µg of GST and the GST-tagged BBS3 and its variants were mixed with 100 µg of bacterially expressed ARL13, ARL13∆N15, ARL13G73L, and ARL13T33N (see above). After incubated for 2 hrs at room temperature, mixtures were purified with GST SpinTrap™ Purification Module (GE Healthcare). Ten micrograms of proteins from elutes was loaded on 12% SDS-PAGE gels for electrogenesis. Gels were then visualized with Coomassie blue staining.

**Size exclusion chromatography (SEC)**

To determine if BBS3 and ARL13 bind each other and the molecule ratio of BBS3 to ARL13 in the transiently existing $BBS3^{GTP}$/ARL13 complex *in vivo*, cilia was prepared from 2.5 liters of *bbs8; BBS3A73L:YFP-TG* cells in the presence of GTPγS (20 mM) or GDP (20 mM) as described above and was equilibrated in 2M NaCl, HEPES (pH 7.0). The ciliary solution was then fractionated on a HiLoad Superdex 16/600 200 pg column (GE Healthcare). To determine the molecule ratio of BBS3 to ARL13 *in vitro*, GST-BBS3A73L/ARL13G73L and GST-BBS3A73L/ARL13T33N complexes obtained as described above was equilibrated in 2M NaCl, HEPES (pH 7.0) and fractionated on a HiLoad Superdex 16/600 200 pg column (GE Healthcare). For both *in vitro* and *in vivo* assays, a total of 180 fractions (1 ml/fraction) were collected. 20 µl of each of 13 fractions (54 to 67 ml for *in vitro* assays and 57 to 70 ml for *in vivo* assays) was analyzed by immunoblotting as described above. Protein standards (Blue Dextran (void volume), Aldolase (158 kD), Ovalbumin (44 kD), Ribonuclease A (13.7 kD), and Aprotinin (6.5 kD) were run on the same column equilibrated in 2M NaCl, HEPES (pH 7.0) to calculate the approximate size of complexes.

***In vitro* GEF assay**

*In vitro* GEF assay was performed to determine GEF activity of BBS3 to ARL13 and GEF activities of IFT25/27 to BBS3 according to the previously described protocol (9). Tandem purification of GST-IFT25/6×His-IFT27, GST-IFT25/6×His-IFT27S79L, and GST-IFT25/6×His-IFT27S30N in bacteria has been described previously (7). GTP affinities of bacterially expressed small GTPases were determined by fluorescence spectrophotometric measurements of 2'(3')-O-(N-methylanthraniloyl)-labeled (mant-labeled) GTP (mant-GTP) (Abcam). To determine mant-GTP binding rate for GTPases, 50 µl of 2× binding reaction buffer (40 mM Tris pH 7.5, 100 mM NaCl, 20 mM $MgCl_2$, and 1.5 µM mant-GTP), 4 µl of each GTPase or mutant alone (50 µM), 36 µl of pure water were added into each assigned well of a 96-well plate (black color to avid light) and mix well by gentle vortex. Time-dependent fluorescence change was recorded once every 30 sec (excitation at 355 nm and emission at 448 nm) using a SynergyH1/H1MF ELISA reader (BioTek). After 5 readings (150 seconds), 10 µl of each of the GEF candidates (5 µM) or pure water was added in wells, mixed well by pipetting, and continued to be read for another 1850 sec. Emission maxima of mant-GTP were plotted against time and normalized by nonlinear regression using GraphPad Prism 8.0 (GraphPad Software). Mant-GTP binding rate was calculated according to the formula: binding rate = Vmax (AFU/sec)/0.75 × Basal mant-GTP AFU × N. Vmax was determined by GraphPad Prism 8.0. Given that the N value for ARL13 is not available yet, we used the N value (N = 2) of CDC42 in all calculations as suggested previously (9).

**Small GTPase assay**

The GTP hydrolysis was measured by an optical assay for the release of inorganic phosphate with the use of reagents from the QuantiChrom™ ATPase/GTPase assay kit (Bioassay Systems) (10). C-terminal 6×His tagged ARL13 and N-terminal GST-tagged BBS3 and BSB3A73Q were expressed in bacteria and used for affinity purification as described above. Free nucleotide was removed with 10 ml G25-resin (GE Healthcare) pre-equilibrated with 20 mM HEPES pH 7.5, 150 mM NaCl. 10 µl (20 µM) GTP-loaded proteins were mixed with 20 µl of 2× assay buffer solutions. For measuring intrinsic or GAP-accelerated GTP hydrolysis, 10 microliters of distilled water and ciliary extracts of *Chlamydomonas* CC-5325 were added. The reactions were performed in 96-well microplates (Corning) at room temperature for 0, 15, 30, 60, and 120 minutes, respectively. After that, 200 µl reagent solutions were added to each reaction and allowed to incubate for 30 min at room temperature. The absorbance at 620 nm was measured with a synergy H1/MF microplate reader (Bio-Tek). Three independently generated experiments were used to calculate the values shown. Preparation of ciliary extracts of *Chlamydomonas* has been described previously (5). The chlorophy11- and GTP-contaminants were removed by applying the extracts through the G25 resin (GE Healthcare). The free phosphate was removed with a Pi-Bind resin (Innova Biosciences).

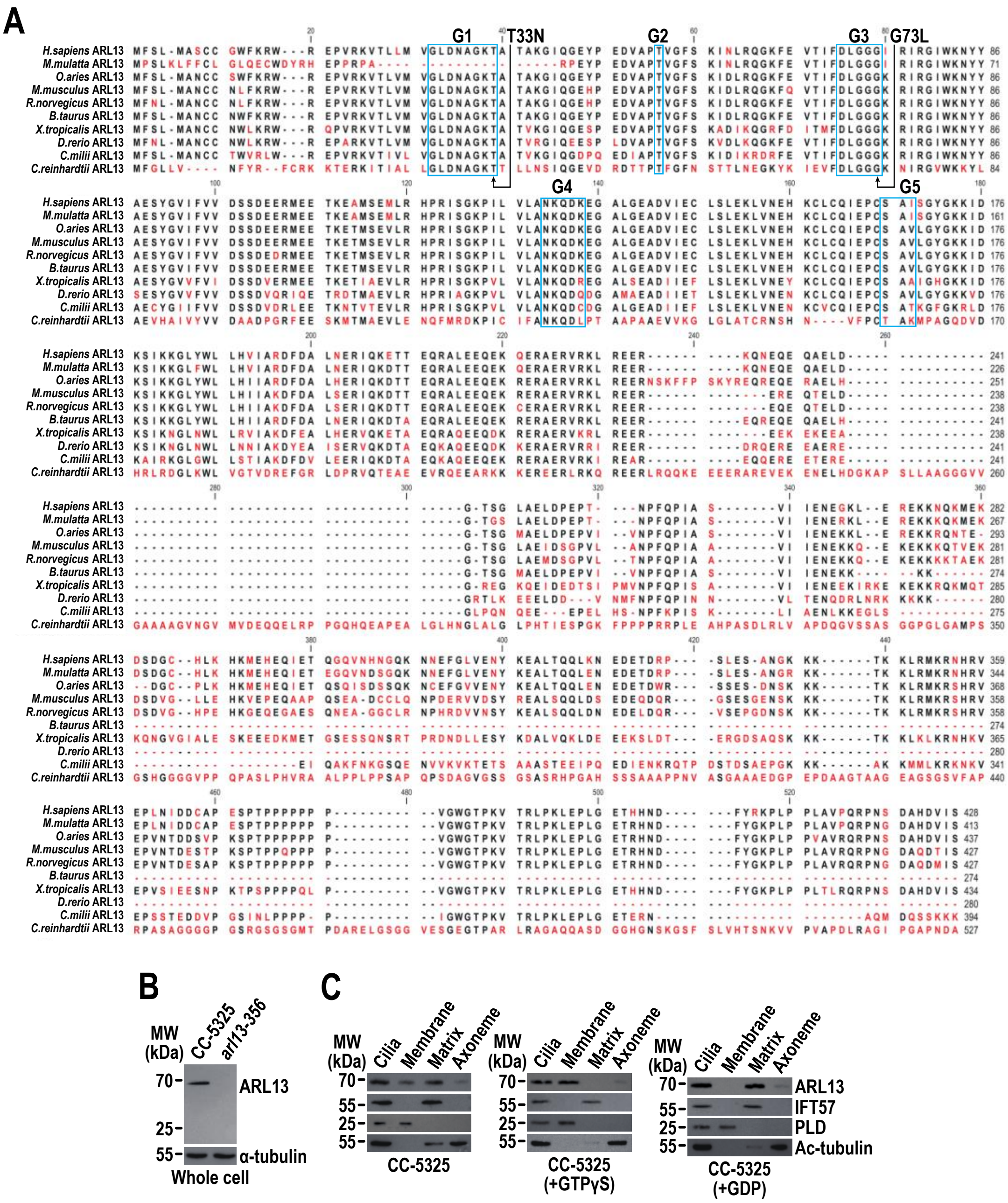


**Fig. S1.** (A) Sequence alignment of deduced amino acid sequences from ten invertebrate and vertebrate ARL13b orthologues. Alignments were generated using CLC main workbench (version 6.8); the most conserved residues are shown in blue, the least conserved are in red. ARL13b contains five conserved domains including G1, G2, G3, G4, and G5 as labeled in the boxes. The P-loop threonine (T33) and glycine (G73) were marked. Dashes indicate gaps introduced to optimize the alignment. Arrowheads indicate missense mutations created in possible dominant-negative and constitutive-active ARL13 mutants. GenBank accession numbers are as follows: *Homo sapiens*, AAI04037.1, *Macaca mulatta*, XP_014984716.2, *Ovis aries*, XP_042086377.1, *Mus musculus*, NP_080853.3, *Rattus norvegicus*, NP_001100571.1, *Bos Taurus*, AAI02961.1, *Xenopus*

*tropicalis*, XP_012811746.1, *Danio rerio,* AAH92689.1, *Callorhinchus milii*, XP_007885211.2 and *Chlamydomonas reinhardtii*, XP_001691430.1. (B) Immunoblots of whole cell sample with the affinity-purified ARL13 antisera identified one band with a size of approximately 70 kDa as expected in CC-5325 rather than the ARL13-null *arl13-356* cells. MW stands for molecular weight. (C) Immunoblots of ciliary fractions of cells indicated on the bottom probed with α-ARL13, α-IFT57 (ciliary matrix marker), α-PLD (ciliary membrane marker) and Ac-tubulin (axoneme marker). MW: molecular weight.

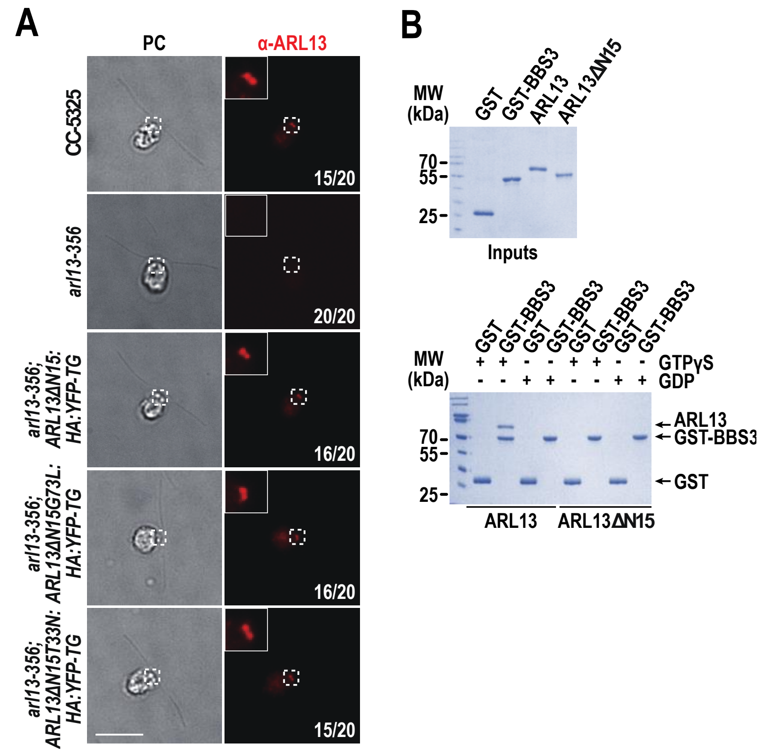


**Fig. S2.** (A) Cells shown on the left stained with α-ARL13 (red). Phase contrast (PC) images of cells were shown. Insets are 9 times enlargements of the basal bodies indicated with a white box. Cell numbers out of 20 cells are listed for representing cells that are ARL13 positive or negative at the basal bodies. Scale bar: 10 µm. (B) GST, GST-BBS3, ARL13, and ARL13△N15 resolved by SDS-PAGE and visualized by Coomassie staining (upper). GST and GST-BBS3 mixed with ARL13 or ARL13△N15 in the presence of GTPγS or GDP and complexes recovered on glutathione beads resolved by SDS-PAGE and visualized by Coomassie staining (bottom). MW stands for molecular weight.

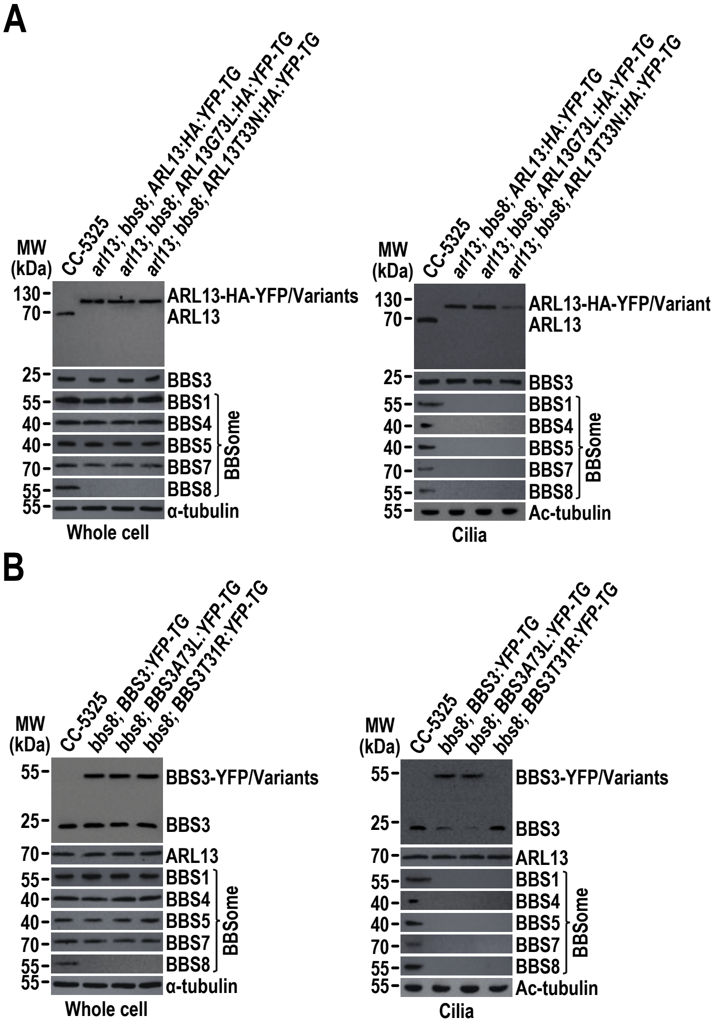


**Fig. S3**. (A-B). Immunoblots of whole cell samples and cilia of cells indicated on the top probed with α-ARL13, α-BBS3, α-BBS1, α-BBS4, α-BBS5, α-BBS7, and α-BBS8. For all panels, α-tubulin and Ac-tubulin were used to adjust the loading for whole cell samples and cilia, respectively. MW: molecular weight.

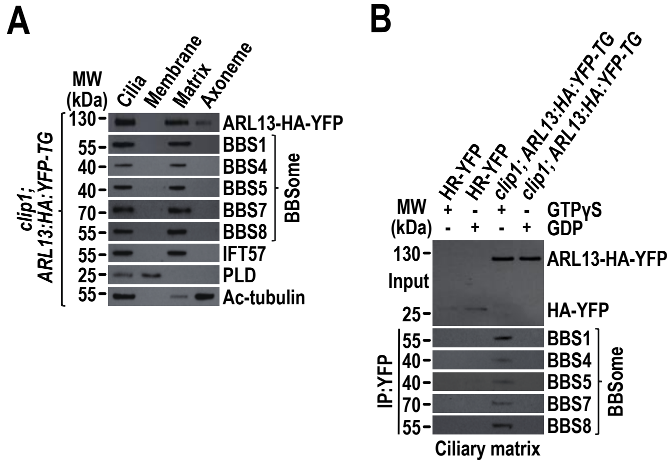


**Fig. S4**. (A) Immunoblots of ciliary fractions of cells indicated on the left probed with α-ARL13, antibodies against the BBSome subunits BBS1, BBS4, BBS5, BBS7, and BBS8, α-IFT57 (ciliary matrix marker), α-PLD (ciliary membrane marker) and Ac-tubulin (axoneme marker). (B) Immunoblots of α-YFP-captured proteins from the ciliary matrix of cells indicated on the top in the presence of GTPγS or GDP probed with antibodies against the BBSome subunits BBS1, BBS4, BBS5, BBS7, and BBS8. Input was quantified with α-YFP by immunoblotting. For both panels, MW stands for molecular weight.

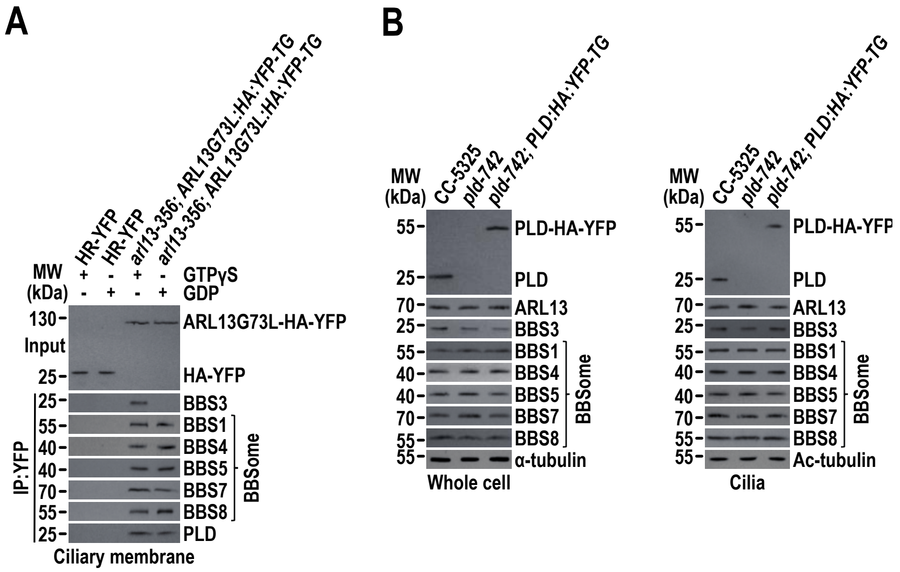


**Fig. S5**. (A) Immunoblots of α-YFP-captured proteins from the ciliary membrane of cells indicated on the top in the presence of GTPγS or GDP probed for BBS3, the BBSome subunits BBS1, BBS4, BBS5, BBS7, and BBS8, and PLD. Input was quantified with α-YFP by immunoblotting. (B) Immunoblots of whole cell samples and cilia of cells indicated on the top probed with α-PLD, α-ARL13, α-BBS3, and antibodies against the BBSome subunits BBS1, BBS4, BBS5, BBS7, and BBS8. Alpha-tubulin and Ac-tubulin were used to adjust the loading for whole cell samples and cilia, respectively. For all panels, MW stands for molecular weight.

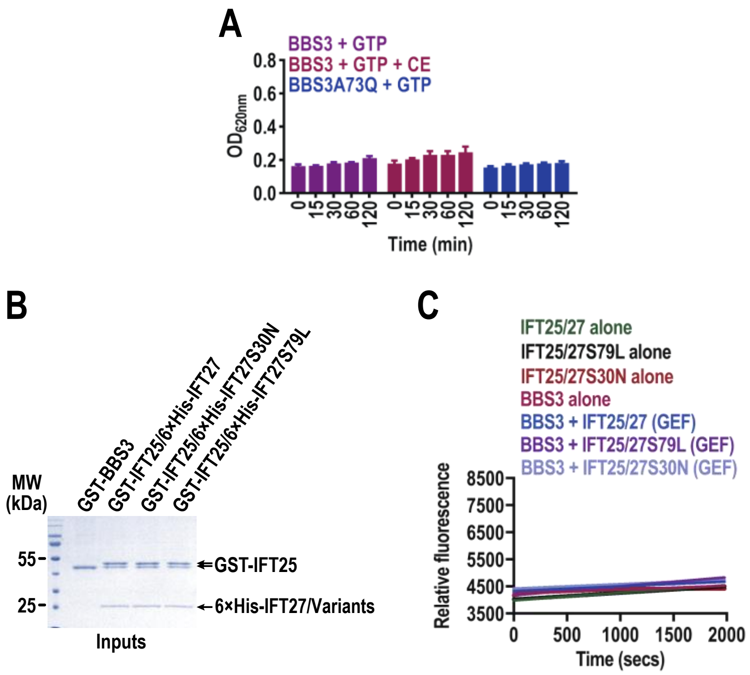


**Fig. S6.** GTP hydrolysis of BBS3 and its A73Q mutant and mant-GTP association measurement for determining GEF activity of IFT25/27, IFT25/27S79L, and IFT25/27S30N on BBS3. (A) GTP hydrolysis of BBS3 and ABBS3A73Q and BBS3 GTP hydrolysis in the presence of CE isolated from CC-5325 cells. BBS3A73Q resembles BBS3 in showing similar but low intrinsic GTPase activity. The presence of CE does not elevate the GTPase activity of BBS3. The assay was performed in a time-course pattern as shown and the inorganic phosphate (Pi) release upon GTP hydrolysis is measured by absorbance at 620 nm. The background absorbance was measured using Pi-free buffer alone or Pi-free buffer plus the purified cell extracts. (B) GST-BBS3 and the tandemly purified GST-IFT25/6×His-IFT27 (IFT25/27), GST-IFT25/6×His-IFT27S30N (IFT25/27S30N), and GST-IFT25/6×His-IFT27S79L (IFT25/27S79L) resolved by SDS-PAGE and visualized by Coomassie staining. Phosphorylated and unphosphorylated GST-IFT25 are shown (8). MW stands for molecular weight. (C) Fluorescence intensity of mant-GTP (0.75 µm) was plotted against the recording time (s) for a combination of 2 µm IFT25/27 and BBS3 alone or in the presence of 0.5 µm IFT25/27, IFT25/27S79L, and IFT25/27S30N (GEF activity on BBS3). Data are averaged from three replicates. The experiment was performed three times, and one representative is shown.

**Table S1.** Antibodies used in this study

| Antibody | Dilution | | Origins | Reference or source |
|---|---|---|---|---|
| | IB | IS | | |
| Anti-ARL13 | 1:250 | 1:200 | Rb | This study |
| Anti-IFT22 | 1:1,000 | N/A | Rb | (3) |
| Anti-IFT38 | 1:1,000 | N/A | Rb | (6) |
| Anti-IFT46 | 1:1,000 | N/A | Rb | (7) |
| Anti-IFT57 | 1:500 | N/A | Rb | (7) |
| Anti-IFT43 | 1:500 | N/A | Rb | (11) |
| Anti-IFT70 | 1:1,000 | N/A | Rb | (7) |
| Anti-IFT139 | 1:1,000 | N/A | Rb | (7) |
| Anti-BBS1 | 1:1,000 | N/A | Rb | (3) |
| Anti-BBS3 | 1:500 | 1:100 | Rb | (3) |
| Anti-BBS4 | 1:500 | N/A | Rb | (2) |
| Anti-BBS5 | 1:1,000 | N/A | Rb | (3) |
| Anti-BBS7 | 1:500 | N/A | Rb | (2) |
| Anti-BBS8 | 1:500 | 1:50 | Rb | (6) |
| Anti-PLD | 1:1,000 | 1:25 | Rb | (2) |
| Anti-α-tubulin | 1:10,000 | N/A | Mo | Sigma-Aldrich (T5168) |
| Anti-acetylated-tubulin | 1:10,000 | N/A | Mo | Sigma-Aldrich (T7451) |
| Anti-GFP (YFP) | 1:1,000 | N/A | Mo | Roche (11814460001) |
| HRP-conjugated goat anti-rabbit IgG | 1:10,000 | N/A | Gt | The Jackson Lab. (111035003) |
| HRP-conjugated goat anti-mouse IgG | 1:10,000 | N/A | Gt | The Jackson Lab. (115035003) |
| Alexa-Fluor 594-conjugated goat anti-rabbit IgG | N/A | 1:400 | Gt | Molecular Probes (A11012) |

Note: Rb, rabbit; Mo, mouse; Gt, goat; HRP, horseradish peroxidase; IB, immunoblotting; IS, immunostaining.

**Table S2.** *Chlamydomonas* strains used in this study

| Name | Genotype | Reference or source |
|---|---|---|
| CC-5325 | *cw15*; $mt^{-}$ | CLiP (12) |
| *arl13-356* | *cw15; arl13::aphVIII*; $mt^{-}$ | CLiP (12) (LMJ.RY0402.181356) |
| *arl13-356; ARL13:HA:YFP-TG* | *cw15; arl13::aphVIII*; $mt^{-}$; *ARL13:HA:YFP-TG* | This study |
| *arl13-356; ARL13G73L:HA:YFP-TG* | *cw15; arl13::aphVIII*; $mt^{-}$; *ARL13G73L:HA:YFP-TG* | This study |
| *arl13-356; ARL13T33N:HA:YFP-TG* | *cw15; arl13::aphVIII*; $mt^{-}$; *ARL13T33N:HA:YFP-TG* | This study |
| *arl13-356; ARL13ΔN15:HA:YFP-TG* | *cw15; arl13::aphVIII*; $mt^{-}$; *ARL13ΔN15:HA:YFP-TG* | This study |
| *arl13-356; ARL13ΔN15G73L:HA:YFP-TG* | *cw15; arl13::aphVIII*; $mt^{-}$; *ARL13ΔN15G73L:HA:YFP-TG* | This study |
| *arl13-356; ARL13ΔN15T33N:HA:YFP-TG* | *cw15; arl13::aphVIII*; $mt^{-}$; *ARL13ΔN15T33N:HA:YFP-TG* | This study |
| *arl13-356*; $arl6^{KD}$; *ARL13:HA:YFP-TG* | *cw15; arl13::aphVIII*; $mt^{-}$; $arl6^{miRNA}$;*ARL13:HA:YFP-TG* | This study |
| *arl13-356*; $arl6^{KD}$; *ARL13G73L:HA:YFP-TG* | *cw15; arl13::aphVIII*; $mt^{-}$; $arl6^{miRNA}$; *ARL13G73L:HA:YFP-TG* | This study |
| *arl13-356*; $arl6^{KD}$; *ARL13T33N:HA:YFP-TG* | *cw15; arl13::aphVIII*; $mt^{-}$; $arl6^{miRNA}$; *ARL13T33N:HA:YFP-TG* | This study |
| *arl13-356; BBS3:YFP-TG* | *cw15; arl13::aphVIII*; $mt^{-}$; *BBS3:YFP-TG* | This study |
| *arl13-356; BBS3A73L:YFP-TG* | *cw15; arl13::aphVIII*; $mt^{-}$; *BBS3A73L:YFP-TG* | This study |
| *arl13-356; BBS3T31R:YFP-TG* | *cw15; arl13::aphVIII*; $mt^{-}$; *BBS3T31R:YFP-TG* | This study |
| *clip1* | *cw15; bbs3::aphVIII*; $mt^{-}$ | CLiP (12) (LMJ.RY0402.149010) |
| *clip1; ARL13:HA:YFP-TG* | *cw15; bbs3::aphVIII*; $mt^{-}$; *ARL13:HA:YFP-TG* | This study |
| *clip1; ARL13G73L:HA:YFP-TG* | *cw15; bbs3::aphVIII*; $mt^{-}$; *ARL13G73L:HA:YFP-TG* | This study |
| *clip1; ARL13T33N:HA:YFP-TG* | *cw15; bbs3::aphVIII*; $mt^{-}$; *ARL13T33N:HA:YFP-TG* | This study |
| *bbs8* | *bbs8*; $mt^{+}$ | (13) |
| *bbs8; BBS8:YFP* | $mt^{+}$; *bbs8;BBS8:YFP-TG* | (1) |
| *arl13; bbs8* | *cw15; arl13::aphVIII; bbs8;* | This study |
| *arl13; bbs8; ARL13:HA:YFP-TG* | *cw15; arl13::aphVIII; bbs8; ARL13:HA:YFP-TG* | This study |
| *arl13; bbs8: ARL13G73L:HA:YFP-TG* | *cw15; arl13::aphVIII; bbs8; ARL13G73L:HA:YFP-TG* | This study |
| *arl13; bbs8: ARL13T33N:HA:YFP-TG* | *cw15; arl13::aphVIII; bbs8; ARL13T33N:HA:YFP-TG* | This study |
| *bbs8; BBS3:YFP-TG* | *bbs8*; $mt^{+}$; *BBS3:YFP-TG* | This study |
| *bbs8; BBS3A73L:YFP-TG* | *bbs8*; $mt^{+}$; *BBS3A73L:YFP-TG* | This study |
| *bbs8; BBS3T31R:YFP-TG* | *bbs8*; $mt^{+}$; *BBS3T31R:YFP-TG* | This study |
| *pld-742* | *cw15; pld::aphVIII*; $mt^{-}$ | CLiP (12) (LMJ.RY0402.071742) |
| *pld-742; PLD:HA:YFP-TG* | *cw15; pld::aphVIII*; $mt^{-}$; *PLD:HA:YFP-TG* | This study |
| *HR:YFP-TG* | *nit1; nit2*; $mt^{+}$; *HA:YFP-TG* | (14) |

Note: CRC stands for *Chlamydomonas* Resource Center.

**Table S3.** Primers used in this study

| Name | Nucleotide sequence |
|---|---|
| **Primers used to amplify the *aph*VIII gene insertion in ARL13 genomic DNA** | |
| gARL13-FOR | 5'-GATTACCATAGCGTTGCTA-3' |
| gARL13-REV | 5'-GTCCCGCCTG GTTCGAGCTC-3' |
| **Primers used to clone target genes** | |
| gARL13ΔN15-FOR | 5'-CCGGATCCATGACCGAGCGAAAGATTAC-3' |
| gARL13ΔN15-REV | 5'-CCGGATCCTGGGTTGAATATGTC-3' |
| gPLD-FOR | 5'-CCTCTAGAATCATCAGGAGCACCTTG-3' |
| gPLD-REV | 5'-CCGAATTCCTTGAACATGTCCCAGAG-3' |
| cARL13-FOR | 5'-GGGAATTCATGTTTGGTTTACTGG-3' |
| cARL13-REV | 5'-CGCTCGAGCTATGCGTCATTGGGG-3' |
| cARL13ΔN15-FOR | 5'-GGGAATTCATGACCGAGCGAAAGATTAC-3' |
| **Primers used to do site-directed mutagenesis** | |
| ARL13G73L-FOR | 5'-GTTTTCGACCTAGGTGGTCTGAAAAACATTC-3' |
| ARL13G73L-REV | 5'-CAGACCACCTAGGTCGAAAACTTCAATCTTG-3' |
| ARL13T33N-FOR | 5'-CTGGATAATGCAGGCAAGAACACTCTGCTAA-3' |
| ARL13T33N-REV | 5'-GTTCTTGCCTGCATTATCCAGGCCTAGCAAC-3' |
| BBS3A73Q-FOR | 5'-CATGTCCGGCCAGGGGAGATACCGGAC-3' |
| BBS3A73Q-REV | 5'-TGGCCGGACATGTCGAACACCGTGAAG-3' |

**Movie S1.** TIRF imaging of ARL13-HA-YFP movement in *arl13-356; ARL13:HA:YFP-TG* cilia. A frame from this movie and kymograph are shown in Figure 1E. Play speed is real-time (15 fps).

**Movie S2.** TIRF imaging of ARL13G73L-HA-YFP movement in *arl13-356; ARL13G73L:HA:YFP-TG* cilia. A frame from this movie and kymograph are shown in Figure 1E. Play speed is real-time (15 fps).

**Movie S3.** TIRF imaging of ARL13T33N-HA-YFP movement in *arl13-356; ARL13T33N:HA:YFP-TG* cilia. A frame from this movie and kymograph are shown in Figure 1E. Play speed is real-time (15 fps).
